# The oncogenic fusion protein EWS-FLI1 promotes premature ageing of biomolecular condensates by catalyzing fibril formation

**DOI:** 10.1101/2022.06.04.494830

**Authors:** Emily E. Selig, Alma K. Romero-Moreno, Shivani Akula, Xiaoping Xu, David S. Libich

## Abstract

Ewing sarcoma (EwS) is an aggressive pediatric cancer of bone and soft tissue. A chromosomal translocation that joins the low-complexity domain of EWS (EWS^LCD^) with the DNA-binding domain of FLI1 (FLI1^DBD^) creates EWS-FLI1, a fusion oncoprotein essential for EwS development and accounts for 85% of all EwS cases. EWS-FLI1 acts as an aberrant transcription factor and interferes with the normal functions of nucleic acid-binding proteins via multivalent interactions and biomolecular condensation. The FLI1^DBD^ was found to directly interact with the EWS^LCD^ causing enhanced phase separation and induced hardening of EWS^LCD^ condensates. Three related ETS DBDs (ERG, ETV1 and PU.1) also induced EWS^LCD^ condensate hardening. DNA binding blocked the interaction with the EWS^LCD^, and NMR spectroscopy confirmed that ETS DBDs interact with EWS^LCD^ via the DNA-binding interface. Our results provide a physical basis for the dominant-negative effect EWS-FLI1 exerts on EWS and highlight the need for further investigations of the FLI1^DBD^-EWS^LCD^ interaction in vivo.

## Introduction

The oncogenic EWS-FLI1 fusion protein is the archetypical example of a related group of fusion proteins characteristic of Ewing sarcoma (EwS), an aggressive pediatric bone and soft tissue cancer^1^. Arising from the t(11;22)(q12;q24) chromosomal translocation that fuses the N-terminal low-complexity domain (LCD) of the RNA-binding protein EWS (EWS) in frame with the DNA-binding domain (DBD) of the E-twenty-six transformation-specific (ETS) family transcription factor Friend leukemia integration 1 (FLI1), the resultant EWS-FLI1 fusion is responsible for approximately 85% of all EwS tumors^1^ (Fig. 1a). EWS-FLI1 acts as a pioneering factor aiding in chromatin opening, yet these observations do not fully explain its oncogenic role in EwS. There is mounting evidence that EWS-FLI1 exerts a dominant negative effect on the normal roles of EWS in transcriptional regulation and splicing^2–5^. Indeed, it was recently noted that the presence of EWS-FLI1 at transcriptionally active sites prevents the release of Breast cancer type 1 susceptibility protein (BRCA1) from DNA-directed RNA polymerase II subunit RPB1 (RNA Pol II), resulting in elevated transcriptional stress and subsequent accumulation of unresolved R-loops^2^.

**Figure 1.**
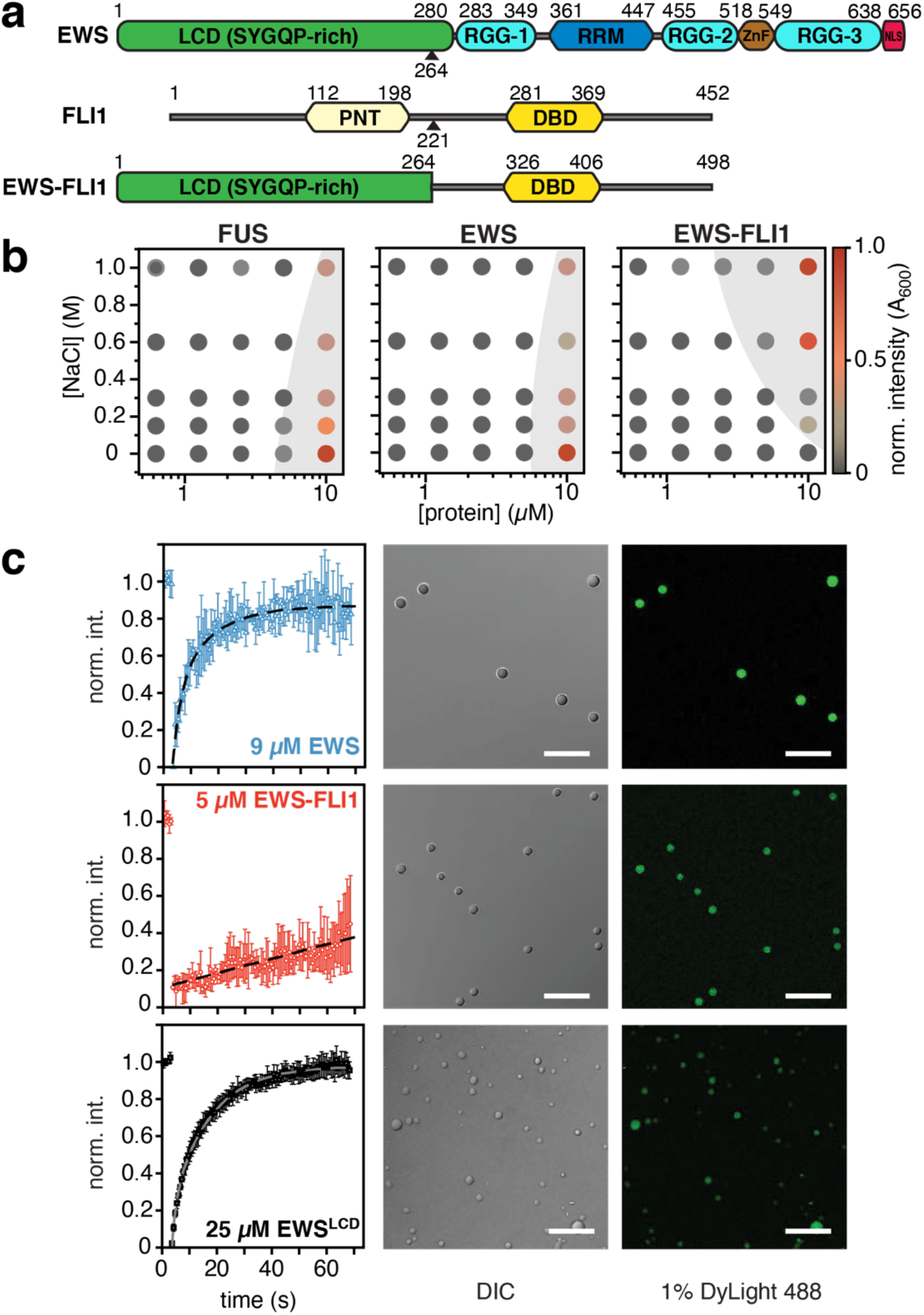
Biomolecular condensation of EWS and EWS-FLI1. **a** The domain architecture of EWS, FLI1 and the EWS-FLI1 fusion protein. The breakpoint for the fusion protein is indicated by a black triangle for EWS and FLI1. **b** Phase diagrams for FUS, EWS and EWS-FLI1 as a function of protein concentration and NaCl concentration. Phase separation was assessed using turbidity measurements at 600 nm. **c** Fluorescence recovery after photobleaching of EWS^LCD 488^ within condensates containing 25 μM EWS^LCD^ with 9 μM WT EWS (blue), 5μM EWS-FLI1 (red) or 25 μM EWS^LCD^ (black). Scale bars indicate 25 μm. FRAP data are presented as mean values ± SEM.

The FET family of RNA-binding proteins is named after its three members: fused-in-sarcoma (FUS), EWS, and TATA-binding protein associated factor 2N (TAF15). All FET proteins are implicated in neurodegenerative diseases amyotrophic lateral sclerosis and frontotemporal dementia, and various cancers including prostate, leukemia and sarcoma^6–9^. FET proteins contain an N-terminal LCD (also known as the transactivation domain) and a C-terminal RNA binding domain (Fig. 1a, Supplementary Fig. 1). The LCD has low sequence complexity, is characterized by a degenerate repeated motif, SYGQ, and lacks stable secondary structure. These properties confer a propensity for self-association and liquidliquid phase separation (LLPS)^10–12^.

The ETS transcription factor family has 28 members characterized by a highly conserved DBD with a winged helix-turn-helix fold that recognizes an eight nucleotide consensus with a core GGAA/T sequence^13^. ETS transcription factors regulate genes involved in processes that can be tumorigenic when dysregulated such as cell cycle control, proliferation, migration, invasion, apoptosis, and angiogenesis^14–17^ For reasons that are not entirely clear, FLI1, transcriptional regulator ERG (ERG), and protein FEV (FEV) most commonly participate in the FET-ETS fusions characteristic of EwS. However, rare chromosomal rearrangements involving FET proteins with other ETS transcription factors have been identified in EwS and in the related family of primitive neuroectodermal tumors (PNETs)^18^.

EWS-FLI1 alters normal genetic programs through aberrant DNA binding at GGAA microsatellites and recruitment of chromatin remodelers, epigenetic modifiers, and transcriptional machinery^11,19–21^. The FLI1 portion of the fusion enables binding to both FLI1 consensus sites and GGAA microsatellites in enhancer regions of EWS-FLI1-responsive genes. However, the EWS LCD contributes to GGAA microsatellite binding and both domains are essential for recruitment of the ATP-dependent BRG1/BRM associated factor (BAF) chromatin remodeling complex and for transcriptional activation^19,22–27^. Transcriptional activation is achieved through the recruitment of RNA Pol II via its C-terminal domain (CTD) in a phosphorylation-dependent manner^11,12,20^.

In recent years, biomolecular condensation, a process in which proteins and nucleic acids demix from the aqueous phase via LLPS to form membraneless organelles, has emerged as a sub-cellular organizational paradigm^28–30^. These membraneless compartments regulate essential cellular processes such as transcription^31,32^, splicing^33^, DNA damage repair^34^ and the stress response^35^. Intrinsically disordered proteins or proteins with intrinsically disordered regions (IDRs), including FET family proteins, are crucial components of biomolecular condensates^10^. Indeed, oncogenic fusions of EWS-FLI1 appear to form dynamic yet specific assemblies with other LCD-containing proteins in cells such as EWS^12,36^ and the CTD of RNA Pol II via multivalent interactions mediated by its LCD^37^. Furthermore, self-association stabilizes the binding of EWS-FLI1 to GGAA microsatellites and is required transcriptional activation, but the exact nature of this self-association remains contentious^11,12,19^.

Since the oncogenic function of EWS-FLI1 requires features of both the EWS^LCD^ and the FLI1^DBD^, the interplay between these two domains was investigated. The properties of biomolecular condensates of EWS, EWS^LCD^ in the presence and absence of EWS-FLI1, FLI1^DBD^ and ETS DBDs were investigated with biophysical approaches. Fluorescent recovery after photo bleaching (FRAP) and Thioflavin T (ThT) assays revealed that the dynamics of intermolecular interactions are altered in EWS-FLI1 condensates relative to EWS and EWS^LCD^ condensates. FLI1^DBD^ catalyzed rapid hardening of EWS^LCD^ condensates, an effect that was inhibited by DNA binding, which inhibited colocalization of FLI1^DBD^ to EWS^LCD^ condensates. The rapid ageing effect was a general feature of ETS domains. NMR confirmed that ETS DBDs transiently interact with EWS^LCD^ via two loops involved in DNA binding. Both the charge density and the relative conformation of the loops was inferred to be important for the EWS^LCD^ interaction. These findings provide a structural explanation for the observed dominant-negative effect of EWS-FLI1 as well as help to explain the apparent toxicity of the fusion, even to EwS tumor cells. Considering these data, further structural investigations as to how the interaction between EWS^LCD^ and ETS DBDs might affect intermolecular interactions of FET-ETS fusions are warranted.

## Results

### EWS-FLI1 has altered condensation properties

Phase diagrams for full-length FUS, full-length EWS and full-length EWS-FLI1 were constructed from turbidity measurements. Phase separation of EWS and FUS was promoted at increased protein and low NaCl concentrations, consistent with previous reports for FUS, where it was determined that phase separation is driven by electrostatic interactions between Arg residues in the Arg-Gly-Gly (RGG) motifs and Tyr residues in the LCD^38,39^ (Fig. 1b). Conversely, phase separation of EWS-FLI1 was promoted by high protein and NaCl concentrations (Fig. 1b), suggesting hydrophobic interactions drive condensation. EWS and EWS-FLI1 are known to interact in vivo, likely via multivalent LCD-LCD interactions in biomolecular condensates^36^. As a way of examining these LCD-LCD interactions, biomolecular condensates were prepared by mixing 25 μM EWS^LCD^, which is common to both EWS and EWS-FLI1 (EWS 1-264, Fig. 1a, Table 1) with 9μM wildtype (WT) EWS, 5 μM EWS-FLI1, or 25 μM EWS^LCD^ (Fig. 1c). WT EWS and EWS-FLI1 readily form dual-component condensates with EWS^LCD^, similar to in vivo observations in a EwS cell line A673^40^. Condensate dynamics were assessed by monitoring fluorescence recovery after photobleaching (FRAP) of the fluorescently labelled molecules within the condensates (Fig. 1c). Condensates containing WT EWS and EWS^LCD^ alone showed rapid FRAP, while condensates containing EWS-FLI1 recovered slower (Fig. 1c), indicating that EWS-FLI1 alters the dynamics of EWS^LCD^ molecules within condensates and that EWS-FLI1 condensates are less fluid and more gel-like.

**Table 1.** Recombinant protein constructs of EWS, FUS, EWS-FLI1 and ETS TFs.

| Construct name | Affinity Tag | MW (kDa) | Margins<sup>1</sup> |
| --- | --- | --- | --- |
| EWS (full-length) | 8 x His | 68.5 | EWS 1 – 656 |
| FUS (full-length) | His-MBP | 53.4 | FUS 1 – 526 |
| EWS-FLI1 (type 2, full-length) | 8 x His | 54.3 | EWS 1 – 264 + FLI1 220 – 452 |
| EWS-FLI1 194-445 F385A | 8 x His | 26.0 | EWS 194 – 264 + FLI1 242 – 452 |
| EWS<sup>LCD</sup> | 8 x His | 27.9 | EWS 2 – 264 |
| EWS<sup>RRM</sup> | 8 x His | 9.9 | EWS 359 – 447 |
| EWS<sup>RRM</sup>-RGG2 | 8 x His | 16.3 | EWS 359 – 513 |
| FLI1<sup>DBD</sup> | 8 x His | 14.6 | FLI1 276 – 399 (F362A) |
| FLI1<sup>DBD</sup> C299S S390C | 8 x His | 14.6 | FLI1 276 – 399 (C299S, F362A, S390C) |
| FLI1<sup>DBD</sup> R2L2 | 8 x His | 14.4 | FLI1 276 – 399 ( R337L, R340L, F362A) |
| FLI1<sup>DBD</sup> Δα4 | 8 x His | 10.3 | FLI1 276 – 361 |
| ERG<sup>DBD</sup> | 8 x His | 14.5 | ERG 306 – 429 (F392A) |
| ETV1<sup>DBD</sup> | 8 x His | 15.1 | ETV1 332 – 458 |
| PU.1<sup>DBD</sup> | 8 x His | 12.3 | PU.1 168 – 270 |

<sup>1</sup> Canonical sequences from Uniprot.

### FLI1^DBD^ alters the physical properties of EWS^LCD^ condensates

The FRAP experiments revealed that FLI1^DBD^ may adversely affect EWS condensate properties. The effect of FLI1^DBD^ on the phase separation propensity of EWS^LCD^ was tested using turbidity measurements and microscopy (Fig. 2a). The addition of equimolar concentrations of FLI1^DBD^ (25μM) significantly increased the turbidity of the sample, indicating an increase in phase separation, and dramatically altered the morphology of the condensates as visualized by bright-field microscopy (Fig. 2a). These condensates no longer coalesce and instead appear fusion-defective (Supplementary Movies 1 and 2). To determine whether this effect was specific to FLI1^DBD^, equimolar concentrations of constructs including the RNA-recognition motif (RRM) of EWS (EWS^RRM^ or EWS^RRM-RGG2^, 25μM) were also tested (Fig. 2a, Table 1, Supplementary Fig. 1). The EWS^RRM^ construct induced a slight increase in turbidity and resulted in EWS^LCD^ condensates that appeared more spherical than those formed in the absence of RRM (Fig. 2a). The EWS^RRM-RGG2^ construct induced an increase in turbidity that was less than that observed for FLI1^DBD^, and induced spherical EWS^LCD^ condensates (Fig. 2a). In contrast to FLI1, EWS^LCD^ condensates formed with the EWS^RRM^ and EWS^RRM-RGG2^ constructs retained the ability to fuse (Supplementary Movies 3 and 4). Therefore, FLI1^DBD^ increases the phase separation propensity of EWS^LCD^, has a different effect on the morphology of the condensates compared to the RNA-binding domains that are naturally found in WT EWS, and changes the material properties of the condensates such that they are fusion-defective.

**Figure 2.**
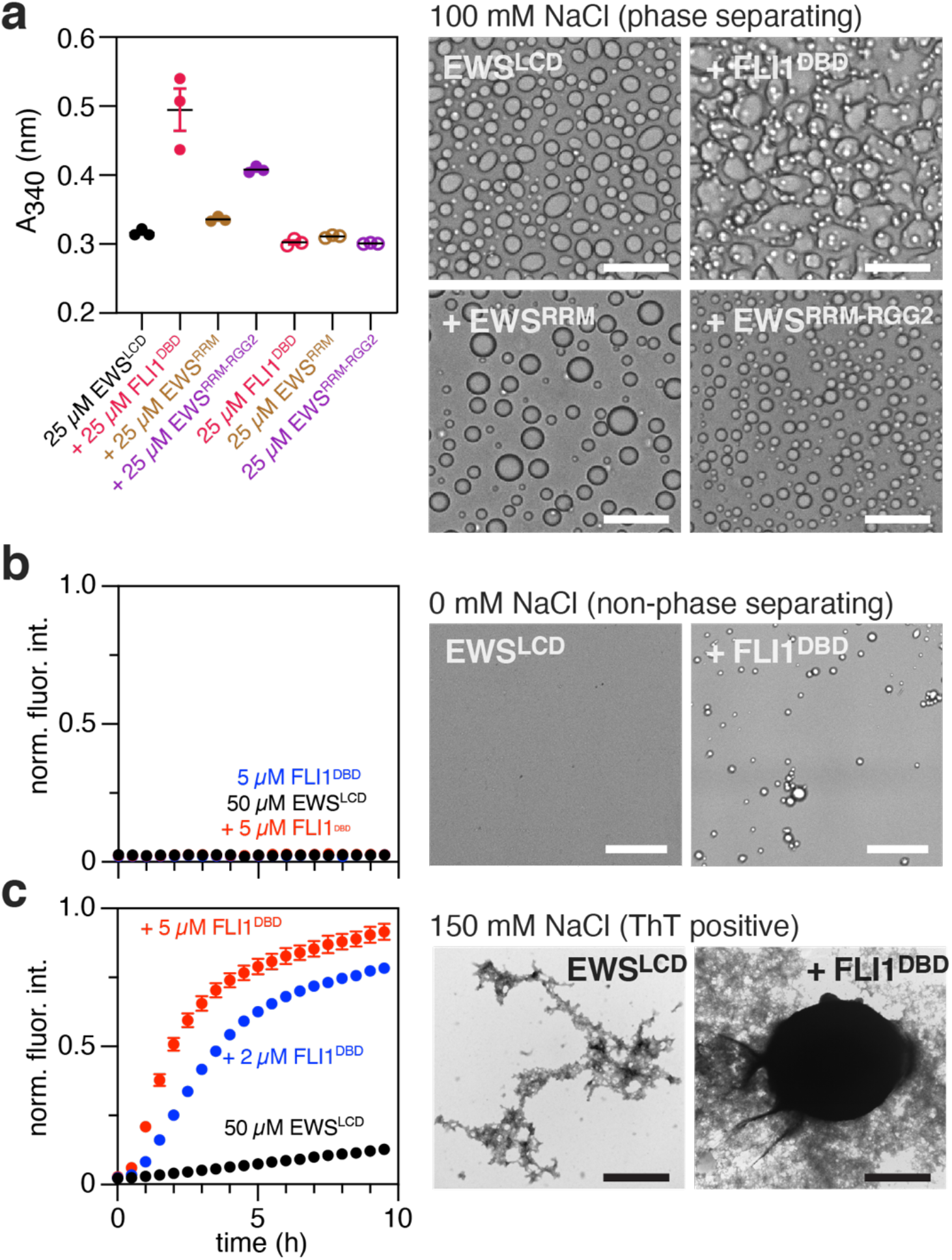
FLI1^DBD^ alters the phase separation propensity of EWS^LCD^. **a** EWS^LCD^ phase separation at 25 μM in 100 mM NaCl assessed by turbidity at 340 nm (left panel) and bright-field microscopy (right panels) alone or in the presence of equimolar concentrations of FLI1^DBD^, EWS^RRM^ or EWS^RRM-RGG2^. Turbidity data are presented as the average of triplicates ± SEM. Scale bars indicate 25 μm. **b** ThT assay of 5 μM FLI1^DBD^ alone, 50 μM EWS^LCD^ alone, and with 5 μM FLI1^DBD^ in non-phase-separating conditions (no NaCl), (left panel) and bright-field images (right panels). Scale bars indicate 50 μm. **c** ThT assay (left panel) of 50 μM EWS^LCD^ alone or with 2 and 5 μM FLI1^DBD^ under phase-separating conditions (150 mM NaCl). Transmission electron micrographs (right panels) of aged samples (T ~ 24 hours) after ThT assays. Scale bars indicate 1 μm. ThT data are presented as mean values ± SEM.

Biomolecular condensates formed by LCD-containing proteins similar in composition to EWS^LCD^ “ripen” or “age” spontaneously over time becoming more gel-like^41^. Cross-β structure has been shown to stabilize the condensed state of the closely related FET protein, FUS^42^, and condensate ageing is concurrent with the formation of fibrillar structures that sometimes protrude from the condensates^41^. To investigate the effect of the FLI1^DBD^ on the formation of cross-β structure in EWS^LCD^ condensates, ThioflavinT (ThT) fluorescence assays were developed. Under non-phase-separating conditions (no NaCl), ThT fluorescence was not observed for EWS^LCD^ alone, or in the presence of FLI1^DBD^ (Fig. 2b). Bright-field microscopy revealed that EWS^LCD^ remains in the dilute phase, but the addition of FLI1^DBD^ was associated with the appearance of several small condensates (Fig. 2b), consistent with FLI1^DBD^ increasing the phase separation propensity of EWS^LCD^ (Fig. 2a), however no measurable increase in turbidity was observed under these conditions. Under phase-separating conditions (150 mM NaCl), EWS^LCD^ condensates age slowly over approximately 10 hours, with a small concomitant increase in ThT fluorescence (Fig. 2c), an effect similar to that reported for FUS^43,44^. The addition of FLI1^DBD^, even at sub-stoichiometric concentrations (1:10 molar ratio), to EWS^LCD^ condensates significantly accelerated condensate ageing in a manner dependent on the concentration of FLI1^DBD^ (Fig. 2c). FLI1^DBD^ also increased ThT fluorescence of WT EWS under phase-separating conditions (Supplementary Fig. 2a). An EWS-FLI1 truncation mutant corresponding to the minimal region of the fusion required for oncogenic transformation^19^ (Table 1, Supplementary Fig. 1) rapidly became ThT positive under phase-separating conditions (Supplementary Fig. 2b).

Aged, ThT positive samples of EWS^LCD^ and EWS^LCD^ with FLI1^DBD^ were analyzed via transmission electron microscopy (TEM) to assess amyloid fibril formation. Although fibrils formed by EWS have been observed^45,46^ and fibril formation within FET protein condensates has also been reported^10,41^, the observed structures did not display the typical morphological features of amyloid fibrils. Instead, heterogeneously branched structures that appeared intertwined were observed that are consistent with a pre-fibrillar or proto-fibril state^45^ (Fig. 2c, Supplementary Fig. 2c). Therefore, the observed increase in ThT fluorescence arises from the formation of cross-β structure that has not had time or is incapable of organizing into bona fide amyloid fibrils. As amyloid fibrils are not known to be associated with EwS, this line of investigation was not pursued and instead ThT fluorescence provided a convenient assay for probing the equilibrium between dilute and condensed phase EWS^LCD^.

### DNA binding to FLI1 inhibits EWS^LCD^ condensate aging

The ability of FLI1^DBD^ to alter EWS^LCD^ condensate properties suggests that the two proteins directly interact. To determine if the interaction site involves the FLI1 DNA-binding site, ThT assays were conducted in the presence a double stranded high-affinity consensus sequence for ETS DBDs^47^ (HA DNA, Table 2). HA DNA inhibited FLI1^DBD^-induced aging of EWS^LCD^ condensates in a concentration-dependent manner (Fig. 3a). At the highest concentrations of HA DNA tested (50 μM corresponding to a 10:1 DNA:FLI1^DBD^ ratio), the effect of FLI1 on EWS^LCD^ condensates was almost entirely abolished (Supplementary Fig. 3a). A double stranded DNA oligonucleotide containing 10 tandem GGAA repeats, which are known to bind FLI1^DBD^ in vitro^27,48^, also inhibited the effect FLI1^DBD^ exerts on EWS^LCD^ condensates (Fig. 3b, Table 2). A scrambled dsDNA control sequence had no inhibitory effect, demonstrating that the inhibitory effect of DNA in these ThT assays is due to DNA binding by FLI1 (Fig. 3b, Table 2).

**Figure 3.**
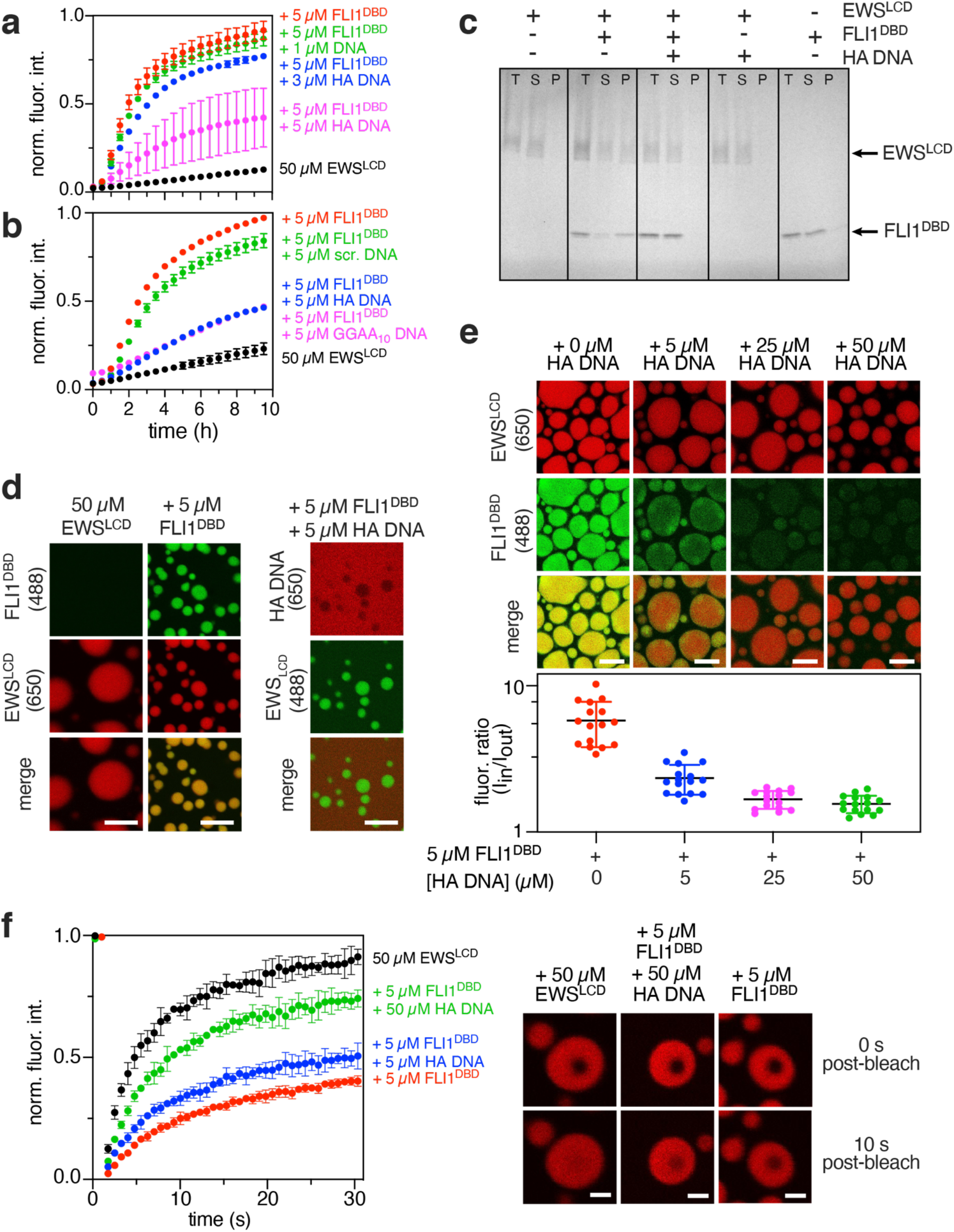
DNA inhibits the effect of FLI1^DBD^ on EWS^LCD^ condensates. Condensates were formed under phase separating conditions (50 μM EWS^LCD^, 150 mM NaCl). **a** ThT assay of 50 μM EWS^LCD^ incubated with 5 μM FLI1^DBD^ (red) and 1 μM (green), 3 μM (blue), and 5 μM (magenta) HA DNA (Table 2). **b** ThT assay of 50 μM EWS^LCD^ alone (black), with 5 μM FLI1^DBD^ (red), or with 5 μM FLI1^DBD^ and 5 μM HA DNA (blue), or GGAA10 DNA (magenta) or scrambled DNA (green) (Table 2). **c** Pelleting assay of aged samples (T ~ 24 hours) of 50 μM EWS^LCD^ incubated alone, with 5 μM FLI1^DBD^, with 5μM of FLI1^DBD^ and HA DNA, or with 5 μM HA DNA. A control sample of 5 μM FLI1 ^DBD^ is also shown. Lanes correspond to the total (T) sample prior to centrifugation, and the supernatant (S) or pellet (P) after centrifugation. **d** Fluorescence and DIC microscopy of 50 μM EWS^LCD^ (1% EWS^LCD 650^) alone (left panels) or with 5 μM FLI1^DBD^ (10% FLI1^DBD 488^) (middle panels). EWS^LCD^ (1% EWS^LCD 488^) condensates with 5 μM unlabeled FLI1^DBD^ and 5μM HA DNA (10% HA DNA^650^) . Scale bars represent 10 μm. **e** Fluorescence microscopy of 50 μM EWS^LCD^ (1% EWS^LCD 650^) with 5 μM FLI1^DBD^ (10% FLI1^DBD 488^) and increasing concentrations of HA DNA. Quantification of the 488 nm-fluorescence intensity ratio Ii⊓/Iout measured in and outside of the condensates. Data are presented as the average of 16 replicates ± SEM. Scale bars indicate 10 μm. **f** FRAP of 50 μM EWS^LCD^ (1% EWS^LCD 650^) condensates alone, with 5μM FLI1^DBD^, or 5 μM FLI1^DBD^ and 50 μM HA DNA. FRAP recovery profiles for 30 seconds post-bleach (right panel) and corresponding images (left panel). Scale bars represent 2 μm. ThT and FRAP data are presented as mean values ± SEM.

**Table 2.**
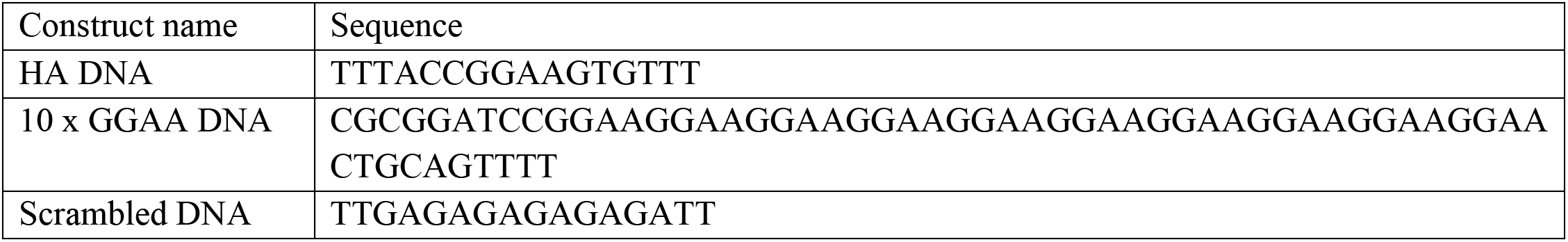
DNA oligonucleotides used for binding studies with ETS DBDs.

To quantify if FLI1^DBD^ associates with EWS^LCD^ condensates or remains in the dilute phase, aged samples (24 hrs old) from ThT assays were pelleted. Though condensates are readily visible via microscopy, the majority (> 95 %) of EWS^LCD^ remains in the dilute phase (supernatant) in the absence of FLI1^DBD^ (Fig. 3c). However, in the presence of FLI1^DBD^, the partitioning of EWS^LCD^ into the pellet fraction was noticeably higher (~ 50 % of total EWS^LCD^, Fig. 3c), consistent with FLI1^DBD^ enhancing EWS^LCD^ phase separation (Fig. 2a). FLI1^DBD^ also partitioned into the pellet fraction (~ 55 % of total FLI1^DBD^), indicating direct association with EWS^LCD^. Addition of HA DNA and FLI1^DBD^ reduced the partitioning of both EWS^LCD^ and FLI1^DBD^ (~ 90 % of each protein remains in the dilute phase) into the pellet further indicating that DNA inhibits the EWS^LCD^-FLI1^DBD^ interaction. HA DNA alone had no effect on EWS^LCD^ phase separation and FLI1^DBD^ incubated alone remained in the dilute phase (Fig. 3c). Furthermore, the partitioning of FLI1^DBD^ with EWS^LCD^ in the pellet was a specific phenomenon as EWS^RRM-RGG2^ remained mostly in the supernatant fraction when incubated with EWS^LCD^ condensates under identical conditions (Supplementary Fig. 3b).

Fluorescence microscopy was used to characterize multicomponent EWS^LCD^ condensates. EWS^LCD,650^ (labeled with DyLight 650) was mixed with FLI1^DBD,488^ (labeled with DyLight 488) under phase-separating conditions. Fluorescence from FLI1^DBD,488^ spatially overlapped with fluorescence arising from EWS^LCD,650^ condensates, consistent with colocalization of FLI1^DBD^ in EWS^LCD^ in condensates (Fig. 3d). FLI1^DBD,488^ does not phase separate under these conditions. This colocalization is specific and further reinforces the notion that EWS^LCD^ and FLI1^DBD^ interact since unrelated proteins such as green fluorescent protein (GFP) do not colocalize to EWS^LCD^ condensates unless fused to EWS^LCD^ (Supplementary Fig. 4a). Surprisingly, mixing EWS^LCD,488^ condensates with unlabeled FLI1DBD and HA DNA^650^ revealed that HA DNA was mostly excluded from the condensates (Fig. 3d).

Exclusion of DNA from EWS^LCD^ condensates may be due to a lack of charge neutralization of the DNA phosphate backbone within the condensates since out of 264 residues the EWS^LCD^ contains only two positively charged residues in contrast with six negatively charged and 27 tyrosines with a delocalized electron in their sidechains. DNA exclusion from the condensate indicated that the inhibitory effect of DNA in the ThT assays may arise from decreased partitioning of FLI1^DBD^ into EWS^LCD^ condensates due DNA binding. To test this, fluorescence microscopy experiments using EWS^LCD,650^, FLI1^DBD,488^, and increasing concentrations of unlabeled HA DNA were conducted (Fig. 3e). A fluorescence intensity ratio (Iin/Iout) was calculated to quantify partitioning of FLI1^DBD^ in the condensates (Fig. 3e). In the absence of DNA, I_in_/I_out_ was ~ 5.8 ± 2.0. As the concentration of HA DNA increased, the intensity of FLI1^DBD,488^ in condensates became noticeably dimmer, and Iin/Iout reduced to 1.6 ± 0.2 for EWS^LCD,650^ with FLI1^DBD,488^ and 50 μM HA DNA (Fig. 3e). Therefore, FLI1 DNA binding out-competes the interactions between EWS^LCD^ and FLI1^DBD^ that drive condensate colocalization. Exclusion of HA DNA from EWS^LCD^ condensates was also observed when full-length EWS-FLI1 was substituted for FLI1^DBD^ (Supplementary Fig. 4b).

FRAP was used to determine whether colocalization of FLI1^DBD^ to EWS^LCD^ condensates results in a reduction in the dynamics of EWS^LCD^ within the condensates, as observed for EWS-FLI1 (Fig. 1c). Freshly prepared EWS^LCD^ condensates underwent rapid recovery after photobleaching, indicating that the condensates are liquid-like (Fig. 3f). In contrast, freshly prepared condensates formed in the presence of FLI1^DBD^ displayed slower fluorescence recovery, indicating that the dynamics of EWS^LCD^ molecules within the condensates change in the presence of FLI1^DBD^ (Fig. 3f). When EWS^LCD^ condensates were prepared with FLI1^DBD^ and increasing concentrations of HA DNA, fluorescence recovered more rapidly in a DNA concentration-dependent manner, consistent with DNA binding inhibiting the effect of FLI1^DBD^ on EWS^LCD^ (Figs. 3e and f).

### The FLI1 DNA-binding and dimer interfaces are not involved in the interaction with EWS^LCD^

The ThT assays and fluorescent colocalization data indicate that FLI1^DBD^ interacts directly with EWS^LCD^. The DNA recognition helix (a3) harbors two highly conserved arginine residues that contact core bases of the ETS consensus sequence in all solved crystal structures of ETS DBDs complexed with DNA^49–53^ (Fig. 4A, Supplementary Fig. 5a). To test if these arginines participate in Arg-Tyr stacking interactions with EWS^LCD^, both arginines were mutated to leucines (R2L2). This mutant is incapable of binding HA DNA (Supplementary Fig. 6a) and its effect on EWS^LCD^ condensates was assessed using ThT assays (Fig. 4b). Surprisingly, the R2L2 mutant retained the ability to enhance the rate of EWS^LCD^ condensate ageing (Fig. 4b). Together these results suggest that the positively charged DNA recognition helix of FLI1 may not be involved in the EWS^LCD^-FLI1^DBD^ interaction.

**Figure 4.**
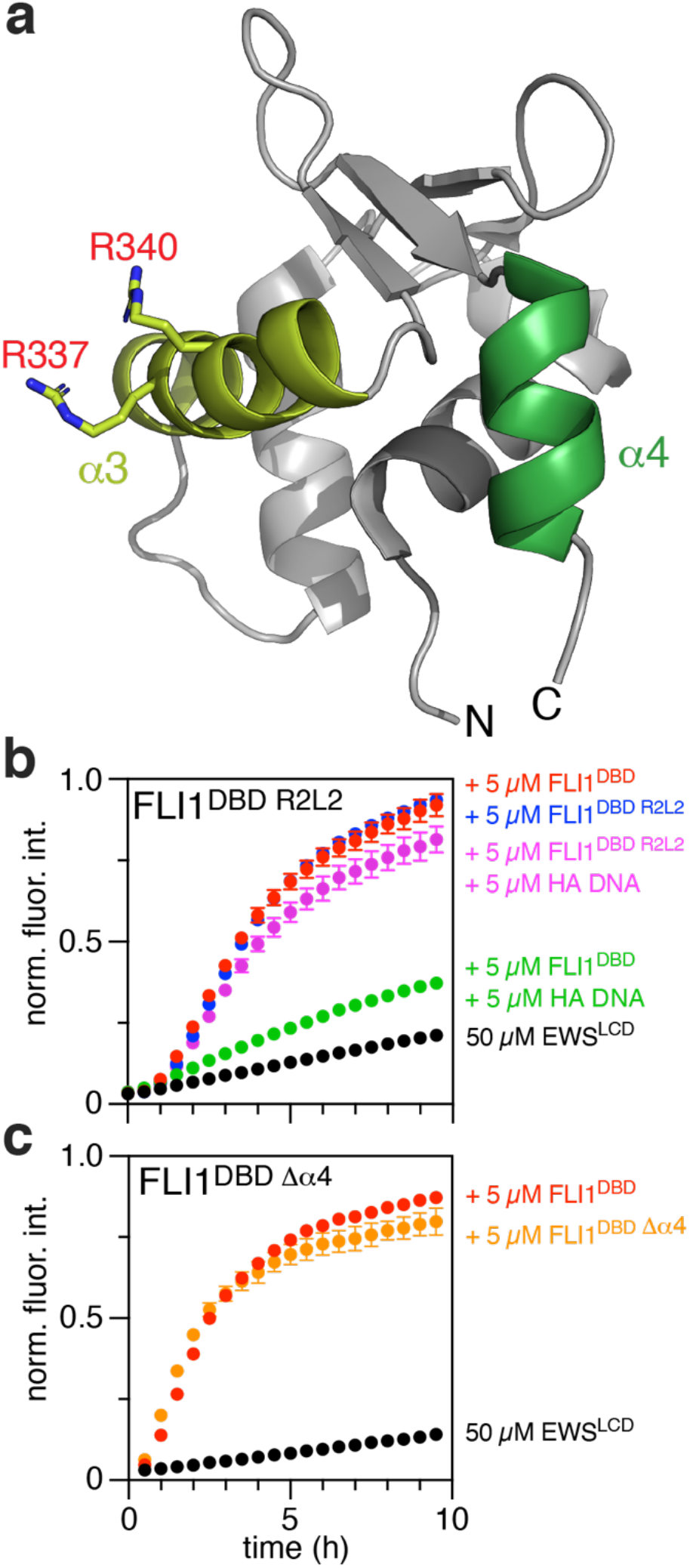
FLI1^DBD^ helices α3 and α4 do not contribute to the ageing of EWS^LCD^ condensates. **a** Cartoon representation of the structure of FLI1^DBD^ indicating the DNA recognition helix (α3), two highly conserved arginine residues that contact DNA (Supplementary Fig. 5), and C-terminal helix α4 (PDB 5e8g). Condensates were formed in the presence of 150 mM NaCl. **b** ThT assay of 50 μM EWS^LCD^ alone (black), or with 5 μM FLI1^DBD^ (red), 5 μM FLI1^DBD^ and 5 μM HA DNA (green), 5 μM FLI1^DBD R2L2^ mutant (blue), or 5 μM FLI1^DBD R2L2^ and 5 μM HA DNA (magenta). **c** ThT assay of 50 μM EWS^LCD^ alone (black), or with 5 μM FLI1^DBD^ (red) or 5 μM FLI1^DBD Δα4^ truncation mutant (orange). ThT data are presented as mean values ± SEM.

The structures of ETS DBDs solved to date share the same fold (Supplementary Fig. 5b), however the precise margins of the ETS domain vary in the existing literature. In some studies an 85 amino acid construct comprising only three α-helices is used^4,54–56^, while other studies used a longer construct with a fourth α-helix^4,22,50^. A recent study crystalized FLI1^DBD^ as a dimer with the fourth α-helix participating in the dimer interface^50^. The FLI1^DBD^ construct used in the ThT assays includes this fourth α-helix (residues 362 - 369) as well as ~30 C-terminal disordered residues. To determine whether this fourth α-helix with exposed hydrophobic residues interacted with EWS^LCD^, a FLI1^DBD^ construct truncated at residue 361was subjected to our ThT assays (Fig. 4a, Table 1, Supplementary Fig. 1). As anticipated, this construct retained DNA binding activity as judged by electrophoretic mobility shift assays (Supplementary Fig. 6b). Further, the shorter FLI1^DBD^ construct exerted the same effect as the longer construct on EWS^LCD^ condensates in ThT assays (Fig. 4c). Together, these observations further delineate the EWS^LCD^ interaction site that is responsible for inducing EWS^LCD^ ageing in condensates to the folded core of FLI1, between residues 276 and 361 but excluding the DNA recognition helix (a3, Fig. 4a).

### Enhancement of EWS^LCD^ condensate ageing is a general property of ETS DBDs

The evidence presented above indicate a specific interaction between EWS^LCD^ and FLI1^DBD^ is responsible for condensate ageing. Asides from FLI1, other EWS-ETS fusions have been identified as driver mutations in EwS. Therefore, two other ETS DBDs that participate in EwS (ERG, ETS translocation variant 1, ETV1), a non-EwS associated ETS DBD transcription factor PU.1 (PU.1), and the EWS^RRM^ and EWS^RRM-RGG2^ control proteins were subjected to ThT assays (Fig. 5, Supplemental Figs. 5b and c). All recombinant constructs were properly folded as judged from circular dichroism spectra (Supplementary Fig. 7). Bright-field microscopy images were acquired for each sample at the end of the ThT assay, (T > 10 hrs) (Fig. 5). The control EWS^RRM^ and EWS^RRM-RGG2^ constructs had minimal effects on the rate at which ThT fluorescence increased for EWS^LCD^ condensates and the condensates remained well dispersed and predominantly spherical (Fig. 5). In contrast, all four ETS DBDs significantly enhanced the rate at which the condensates aged (Fig. 5). Furthermore, EWS^LCD^ condensates formed with ETS DBDs were irregularly shaped, most notably in the presence of PU.1^DBD^ and ETV1^DBD^ (Fig. 5). At higher concentrations of FLI1^DBD^ (> 50 μM), EWS^LCD^ condensates were highly irregularly shaped, even at T ~ 0 hours (Supplementary Fig. 8).

**Figure 5.**
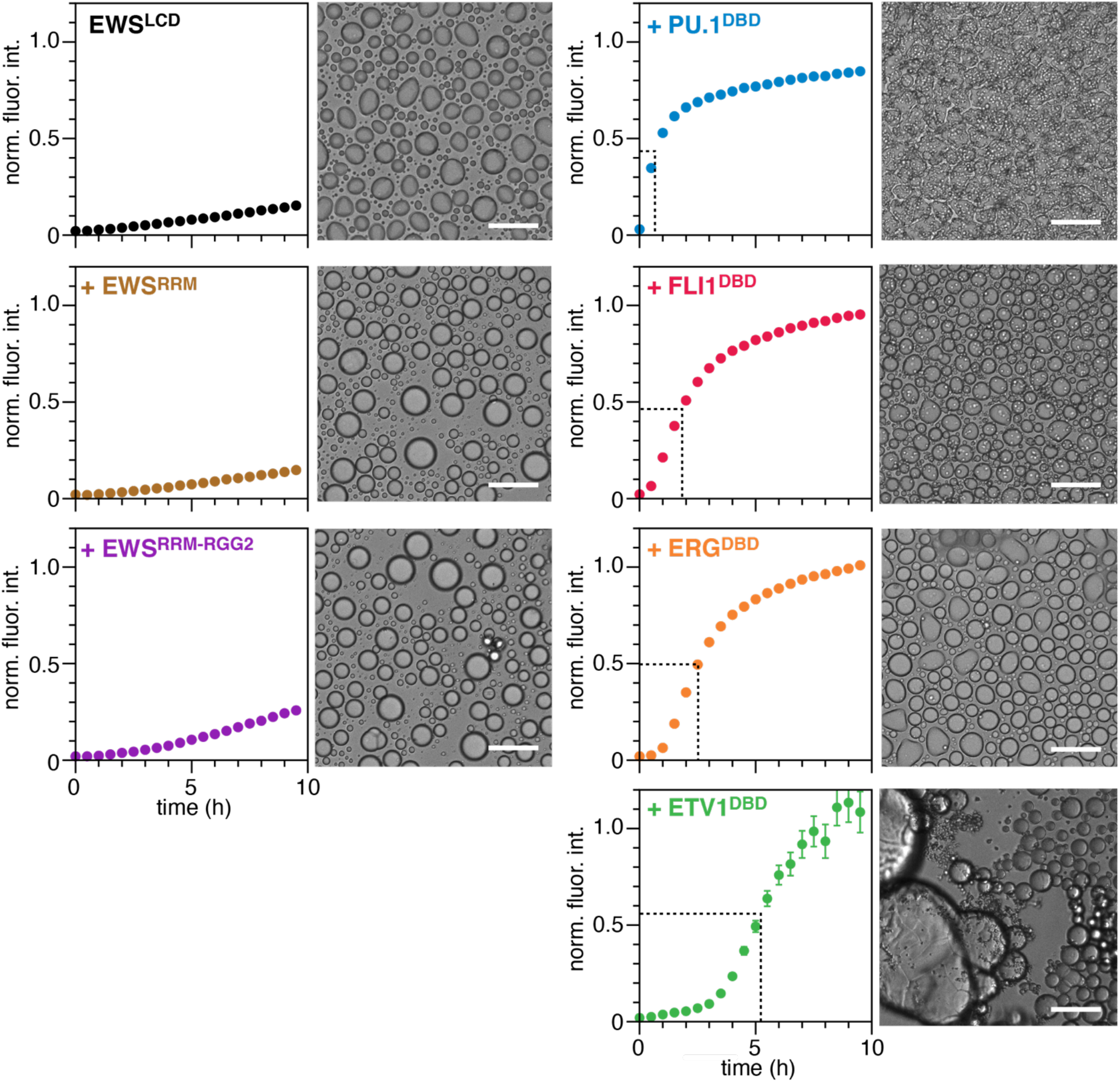
Ageing effect on EWS^LCD^ condensates is conserved for other ETS DBDs. ThT assays (left panels) and bright-field microscopy of aged samples at the end of the ThT assays (T ~ 24 hours, right panels) of 50 μM EWS^LCD^ incubated alone or with 5 μM of EWS^RRM^, EWS^RRM-RGG2^, or PU.1^DBD^, FLI1^DBD^, ERGD^BD^, and ETV1^DBD^ as indicated. All samples were prepared with 150 mM NaCl. Dashed lines estimate half the maximum signal. Scale bars indicate 50 μm. ThT data are presented as mean values ± SEM.

The addition of HA DNA along with ERGD^BD^ and PU.1^DBD^ revealed that DNA binding had the same inhibitory effect on EWS^LCD^ condensate ageing as was observed for FLI1^DBD^, suggesting a common mechanism for ETS domains (Supplementary Fig. 9a). Additionally, the ETS DBD constructs, EWS^RRM^ or EWS^RRM-RGG2^ constructs alone were incapable of forming ThT-positive structures (Supplementary Fig. 9b). Potential contribution of the 8x His-tag used for purification of ETS DBDs, EWS^RRM^ and EWS^RRM-RGG2^ proteins was assessed with a His-tag free version of PU.1^DBD^ (Supplementary Fig. 9c). Since the EWS^RRM^ and EWS^RRM-RGG2^ constructs also retained His-tags but did not induce condensate ageing and the His-tag free PU.1^DBD^ induced condensate ageing at the same rate as His-tagged PU.1^DBD^, the His-tag did not non-specifically induce ageing of EWS^LCD^ condensates (Supplementary Fig. 9c). These finding suggest that the effect FLI1^DBD^ exerts on EWS^LCD^ condensates is conserved for members of the ETS TF family.

### ETS DBDs interact with the EWS^LCD^ via residues adjacent to the DNA-binding face

Solution NMR was used to map the ETS DBD interface with the EWS^LCD^. The interaction between the proteins was predicted to be transient, and significant technical challenges arise from using NMR to study an aggregation-prone system. PU.1^DBD^ was chosen for NMR since it induced the most rapid increase in ThT fluorescence, indicative of a stronger interaction between PU.1^DBD^ and EWS^LCD^ (Fig. 5). Backbone resonances of PU.1 were assigned using standard approaches^57^. ^1^H,^15^N heteronuclear single quantum coherence (HSQC) spectra were recorded for each addition of unlabeled EWS^LCD^ to ^15^N PU.1^DBD^ up to a molar ratio of ~ 3:1 (Supplementary Fig. 10). Addition of higher concentrations of EWS^LCD^ were not possible because, PU.1^DBD^ induced EWS^LCD^ self-association and phase separation. Nevertheless, small chemical shift perturbations (CSP) (δ△ ~ 0.04 ppm) were observed for PU.1 resonances (Fig. 6a, Supplementary Fig. 10). The signal intensity uniformly decreased between the initial and last titration points likely due to co-aggregation of PU.1 with EWS^LCD^ condensates (Fig. 6b). However, a few peaks were differentially broadened, and coincided with or were located near residues with CSPs (Figs. 6a and b). Residues with CSPs greater than one standard deviation and residues with differential signal intensities less than one standard deviation were plotted onto the AlphaFold^58^ structure of human PU.1 (Fig. 6c). Notably, these residues clustered to one face of the DBD, primarily incorporating residues in the ETS DBD “wings” (Loops 4 and 6) that contact the phosphate backbone of DNA^51^, however no shifts or differential broadening were observed for the DNA recognition helix, supporting our earlier hypothesis that it is not involved in the interaction with EWS^LCD^ (Fig. 6d). Furthermore, no shifts or peak broadening were observed on the opposite face of the DBD (Fig. 6c). The CSPs and differential broadening observed for the disordered C-terminal tail were deemed non-specific since they are not conserved in other ETS DBDs and since C-terminally truncated ETS domains (FLI1^DBD,Δα4^) retain condensate ageing activity (Fig. 4c).

**Figure 6.**
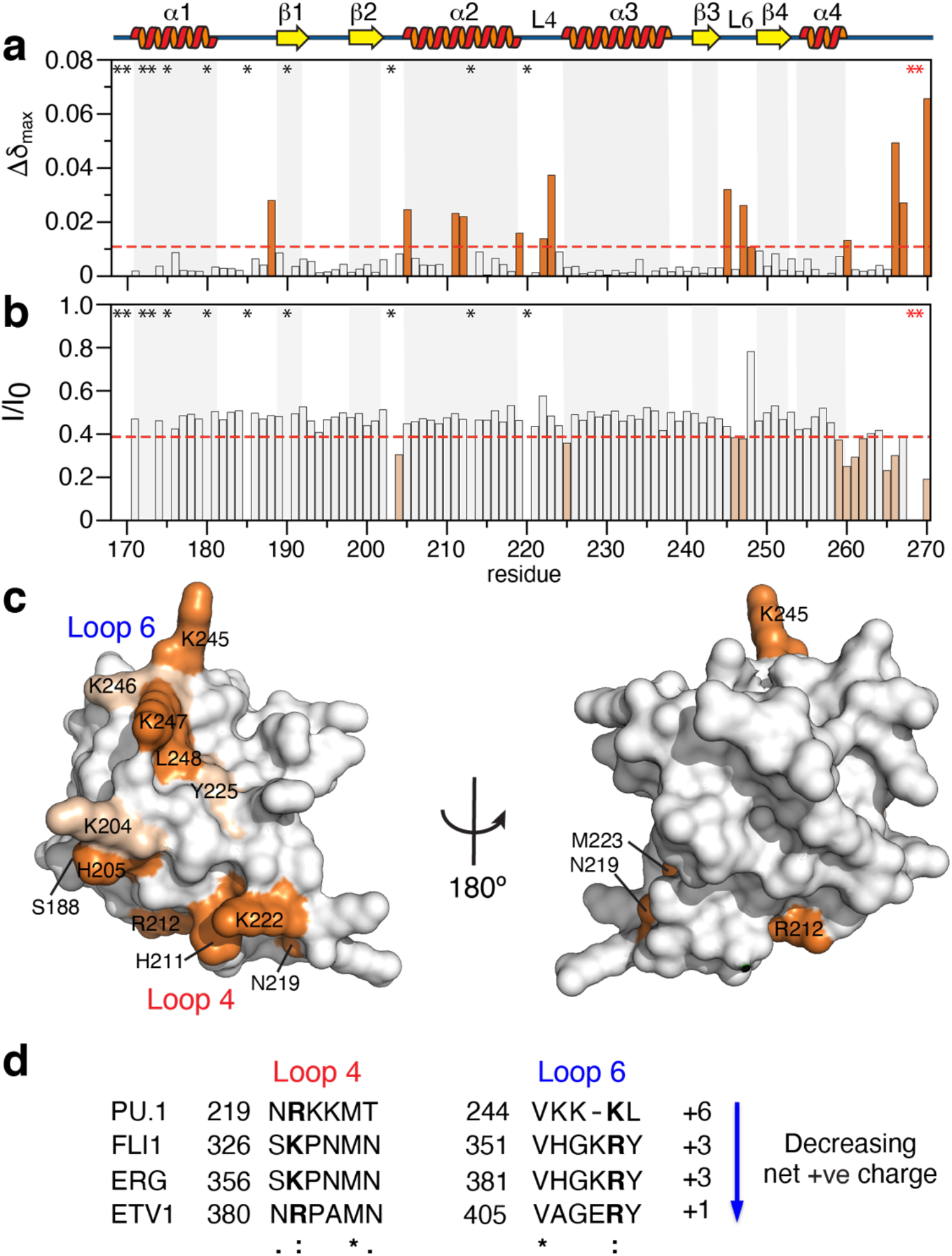
Residues in the DBD wings of PU.1 are involved in the interaction with EWS^LCD^. **a** Chemical shift perturbations and **b** signal broadening of ^15^N-labelled PU.1^DBD^ residues, upon titration with a 3:1 molar ration of EWS^LCD^. Red horizontal dashed lines indicate the standard deviation of the CSP or intensity ratio. Residues with CSPs (intensity differences) greater than 1 standard deviation are plotted in orange. Overlapped and ambiguously assigned residues are indicated by black asterisks and two C-terminal proline residues are denoted by red asterisks in (**a**) and (**b**). **c** Residues with CSP (broadening) greater (less) than the standard deviation in (**a**) and (**b**) were mapped to the AlphaFold structure of the human PU.1^DBD^ (168-257). **d** Sequence alignment of loops 4 and 6 that were identified to interact with the EWS^LCD^. The net charge summed across the two loops is indicated, conserved positive charges in each loop are bolded (Supplementary Fig. 5).

Comparison of the sequences of PU.1 loops 4 and 6 to those of ERG, FLI1 and ETV1 revealed that the total charge varies between +6 to +1 between the DBDs (Fig. 6d). The loops in PU.1 are flexible and enriched in Lys residues and thus have a high net charge of +6, loops 4 and 6 of FLI1 and ERG are identical with a net charge of +3, and ETV1 has a net charge of +1 (Supplementary Fig. 11). Each loop contains at least one highly conserved positively charged residue (Arg or Lys) at 220 and 247 (PU.1 sequence numbering, Fig. 6d). The ThT assays revealed that PU.1^DBD^ induced condensate ageing at the fastest rate, followed by FLI1 ^DBD^ and ERGD^BD^, and ETV1^DBD^ induced ageing at the slowest rate (Fig. 5). Therefore, the net charge of loops 4 and 6 is correlated to the rate at which the ETS DBD induces EWS^LCD^ condensate ageing with higher overall charge inducing the fastest ageing. This correlation suggests that the interaction between ETS DBDs and EWS^LCD^ that drives condensate ageing is influenced by electrostatic interactions. However, electrostatic interactions are clearly not the only factor that influences condensate ageing because the EWS^RRM-RGG2^ construct is enriched with positively charged Arg residues, yet this construct does not affect EWS^LCD^ condensate ageing like the ETS DBDs (Fig. 5). Therefore, it is likely that the relative positioning of the two loops in the ETS DBD structure is also an important factor contributing to EWS^LCD^ condensate ageing.

## Discussion

Though the roles of EWS are not completely defined, knockout of EWS is postnatal lethal in mice, indicating it functions in homologous recombination, the DNA damage response, and in splicing^2,59–61^. The phenotype induced by expression of EWS-FLI1 mimics EWS knockdown phenotype in HeLa and EwS cells leading to hypothesis that EWS-FLI1 exerts a dominant-negative effect on the normal cellular functions of EWS^2,4,5,62^. This work demonstrated that FLI1^DBD^ enhanced the phase separation propensity of EWS^LCD^, localized specifically to EWS^LCD^ condensates, and accelerated condensate ageing. This effect was conserved for three other ETS DBDs. Boulay et al. recently reported that b-isoxazole mediated precipitation of EWS-FLI1 in EwS cell lines was enhanced relative to WT EWS^19^, supporting the hypothesis that the interaction between EWS^LCD^ and FLI1^DBD^ enhances phase separation. Previous studies in prostate cancer cell lines also identified interactions between EWS and ERG, ETV1, ETV4 and ETV5 and these were proposed to be necessary and sufficient for the tumor phenotype^6^. Furthermore, co-immunoprecipitation experiments demonstrated a direct interaction between PU.1 and FUS that inhibits the normal functions of FUS in splicing^63,64^. Therefore, mis-localization of ETS domains either as fusions or due to aberrant regulation negatively impacts the function of EWS and possibly other nucleic acid-binding proteins. The interaction between ETS DBDs and EWS^LCD^ appears to involve residues that are at least partially occluded by DNA binding. As a result, DNA binding reduces the interactions between ETS DBDs and EWS^LCD^ that modulate phase separation and promote the formation of cross β-structure within EWS^LCD^ condensates (Fig. 7). The in vivo situation is more complicated since other proteins, nucleic acids, and post-translational modifications will modulate the properties of the condensate, thus fibril formation is not a likely endpoint in EwS, rather the dominant negative effect is attributable to enhanced self-association of EWS and likely other, related proteins.

**Figure 7.**
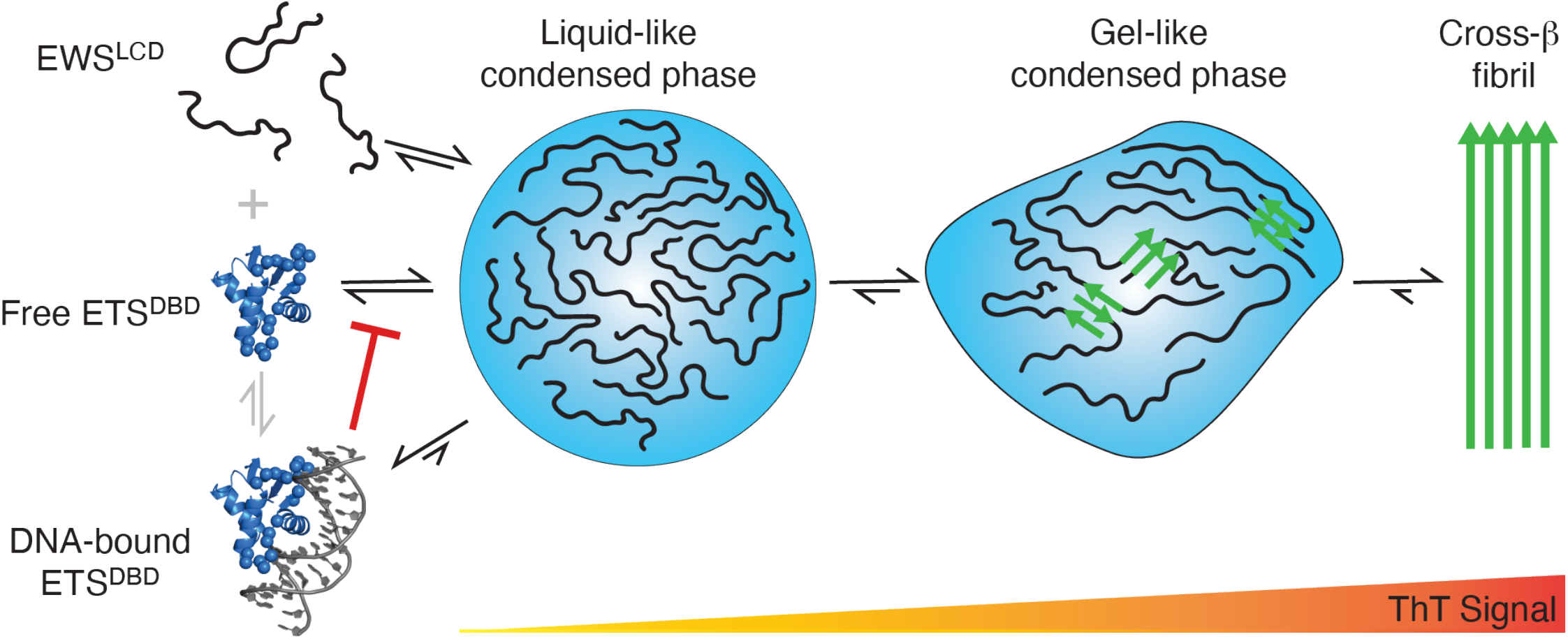
A model for the effect of ETS DBDs on biomolecular condensation by EWS^LCD^. EWS^LCD^ exists in an equilibrium between monomers, liquid-like condensates, gel-like condensates, and amyloid fibrils. Colocalization of ETS DBDs to EWS^LCD^ condensates, enhances the propensity for EWS^LCD^ to phase separate and increases the rate at which the condensates age, forming ThT-positive structures. DNA-binding by free ETS DBDs inhibits colocalization to EWS^LCD^ condensates and thereby reduces the effect ETS DBDs exert on EWS^LCD^ condensates.

Self-association of EWS^LCD^ stabilizes EWS-FLI1 binding to GGAA microsatellites, helps recruit RNA Pol II, and is required for transcriptional activation of GGAA-associated genes that drive oncogenesis^12,19,20^. Chong et al. over expressed exogenous EWS^LCD^ in EwS patient-derived cells to assess the roles of EWS and EWS-FLI1 on transcriptional output of GGAA-responsive genes. Increasing the population of EWS^LCD^ reduced the transcriptional activation by EWS-FLI1 likely through increased LCD-LCD interactions^40^. The authors proposed that a narrow optimum range of LCD-LCD interactions are required to activate transcription. When considered through the model presented here, the observations of Chong et al., raise an interesting conjecture; could excess EWS^LCD^ act as a molecular sponge and blunt the effects of EWS-FLI1 on transcriptional activation? Therefore, the question remains whether the tunable transcriptional activity of EWS-FLI1 arises from the number of molecules participating in the transcriptional hubs or whether it results from tight regulation of the dynamics of intermolecular interactions at these hubs. The tunable LCD-LCD interactions that appear to regulate the transcriptional output of EWS-FLI1 need to be further investigated in situ in the context of the fulllength EWS-FLI1 fusion protein.

EWS^LCD^ will spontaneously form ThT-reactive structures (albeit slowly) and this reaction is much more rapid when catalyzed by ETS DBDs. The critical residues mediating interactions between EWS^LCD^ and PU.1^DBD^ were localized to loops 4 and 6 of the DBD. The overall positive charge of these loops correlated with the rate at which ThT-positive structures formed and therefore positively charged residues, such as lysines, (enriched in PU.1) may contribute to the affinity of ETS DBDs for the EWS^LCD^. Supporting the role of positively charged residues in mediating the interaction with EWS^LCD^, lysine residues in the CTD of RNA Pol II were found to be important for its interaction with hydrogels and condensates formed by the FET family proteins, TAF15 and FUS^38,65^. Despite the observed correlation with positive charge, the conformation of the ETS DBD loops also contribute to EWS^LCD^ aggregation.

While TEM did not conclusively demonstrate the formation of amyloid fibrils, fibril formation by EWS has been observed previously^45,46^, though unlike typical amyloid fibrils, these are labile to disassembly^45,66^. Furthermore, hydrogels of the LCD of FET family proteins have been shown to consist of amyloid-like fibrils^10,11^. There are no high resolution structural models for EWS fibrils, although several models exist for short segments of FUS^67,68^ and for the entire LCD of FUS^42^. Assuming EWS^LCD^ forms cross-β structures in a similar way, via the formation of parallel, in-register β-strands, a key structural feature of these fibrils would be “ladders” of identical sidechains stacked at an interval corresponding to the interstrand distance^69^. These stabilizing ladders could be formed by aromatic tyrosine residues that undergo π-π stacking or by glutamine/asparagine residues that form complimentary hydrogen bonds across the stacked β-strands^69^. Structural insight remains elusive for how ETS DBDs enhance the rate at which EWS^LCD^ condensates become ThT positive, but tyrosine residues in the EWS^LCD^ are important for functional self-association^19,70^. Therefore, tyrosine residues in EWS^LCD^ may interact with the positive charges (cation-π) on ETS DBD loops in a conformation conducive for aromatic ladder formation which in turn may serve to catalyze the formation of cross-β structure. This interaction is expected to be extremely transient and thus the increased high-local concentration of the EWS^LCD^ in condensates enhances the rate of cross-β formation, consistent with the observation that the rate of ThT-positivity is enhanced under phase-separating conditions. The CTD of RNA Pol II has been shown to bind to hydrogels comprised of fibrillar polymers formed by the LCD of FET family proteins, and the degree of CTD binding correlates with the degree of transcriptional activation^11^. It is therefore possible that the interaction between FLI1^DBD^ and EWS^LCD^ in the fusion protein alter the biophysical properties of EWS-FLI1 condensates in a way that enhances recruitment of RNA Pol II via CTD interactions.

While fluorescence microscopy revealed, unexpectedly, that HA DNA is mostly excluded from condensates formed by EWS^LCD^, the in vivo situation involving WT EWS and other proteins is likely to be different. Cation-π interactions between tyrosine residues in the LCD and arginine residues in the RGG motifs likely modulate phase separation of WT EWS, as has been observed for FUS^71^. Technical considerations including preparation of NMR amenable samples and relative instability of the full-length constructs precluded the use of WT EWS and EWS-FLI1 constructs. EWS-FLI1 constructs with solubility tags are less aggregation prone^72^, however even these constructs do not form condensates with rapid fluorescence recovery after photobleaching^20^ and the effects of solubility tags confound study of the self-associative properties of the protein. Although DNA colocalization with EWS-FLI1 containing condensates was not observed in this work, it is plausible that in vivo additional binding partners and post-translational modifications of EWS-FLI1 may serve to neutralize the net negative charge that could be expected of a biomolecular condensate formed by EWS-FLI1 and DNA.

EWS-FLI1 drives oncogenesis through dysregulation of genes downstream of GGAA microsatellites and dysregulation of alternative splicing programs, a function dependent on an intact DBD^1,61,73^. As part of its oncogenic function, EWS-FLI1 also associates with a multitude of nucleic acid binding proteins, including the remaining copy of EWS^36^, FUS, and RNA Pol II^3,21,74^. These interactions remain poorly understood, likely due to their heterogenous and transient nature. The central finding presented here is that ETS DBDs colocalize to and interact transiently yet specifically with the EWS^LCD^, altering the intermolecular interactions that govern its LLPS and subsequent condensate ageing (Fig. 7). These results provide a mechanism that may explain how EWS-FLI1 interferes with the normal functions of EWS and potentially other crucial nucleic acid binding proteins. The presence of the FLI1^DBD^ (or other ETS DBD) promotes self-association and aggregation of EWS altering the local dynamics and intermolecular interactions crucial for EWS function.

## Methods

### Protein expression and purification

Protein constructs used are listed in Table 1. All constructs were derived from human sequences, codon optimized for *E. coli* expression, synthesized (GenScript) and cloned into modified pET expression vectors that placed a 6 or 8 x His-tag followed by a tobacco etch virus (TEV) protease cleavage site N-terminal to the protein coding sequence. The F385A mutation in EWS-FLI1 (EWS-FLI1 numbering) and the F362A mutation in FLI1^DBD^ (FLI1 numbering) correspond to the same residue and were introduced to reduce the propensity of FLI1^DBD^ to dimerize^50^. Full-length MBP-FUS was a gift from N. Fawzi (Addgene plasmid #98651). All plasmids were verified by DNA sequencing. Constructs were expressed and purified from *E. coli* BL21 Star™ (DE3) or BL21 (DE3) pLysS (Invitrogen, MA) cells in Luria Broth or M9 media supplemented with ^15^NH_4_C1 (1 g/L, Sigma) and 0.02% (w/v) yeast extract for ^15^N labelling or ^15^NH_4_C1 and ^13^C6 D-glucose (3 g/L, Sigma) with 0.02% (w/v) Isogro®-^13^C, ^15^N (Sigma) for ^15^N, ^13^C labelling. Generally, expression was induced at OD600 of 0.6 – 0.8 with 1 mM IPTG and continued for 4 hours at 37°C, or overnight at 16°C or 12°C.

All proteins were purified by immobilized metal affinity chromatography or cation exchange chromatography fractionation, followed by cleavage of the His-tag using TEV S219V protease^75^ and a final polishing size-exclusion chromatography (SEC) step. Most ETS DBDs (unless otherwise specified) were purified without cleavage of the His-tag. EWS and FUS constructs were purified in 20 mM CAPS pH 11 as previously described^57,76^. ETS DBDs and EWS RNA binding domains were purified in either sodium phosphate buffers or Tris buffers at pH 7-8. Protein purity for all constructs was confirmed to be ≥ 95% by SDS-PAGE. Protein aliquots were stored at -80 °C (see Supplementary Material for detailed protocols).

### Fluorescent labelling

Purified EWS^LCD^, EWS and EWS-FLI1 were fluorescently labelled with DyLight 488, Dylight 650 or fluorescein maleimide 488 (Thermo Scientific) using sortase^77^. The dyes were resolubilized in DMF and used to label the SortC1 peptide, KLPETGG (GenScript). Proteins to be fluorescently labelled were prepared in 0.1 M sodium bicarbonate pH 9.5 with equal concentrations of the dye-labelled peptide and 2.5 μM sortase, and incubated overnight at room temperature protected from light. Free dye, peptide and enzyme was removed by SEC using a HiLoad 16/600 Superdex 75 pg column (Cytiva), labelling was verified by SDS-PAGE, and UV transillumination. Fluorescently labelled protein was stored at -80°C until use. FLI1^DBD C299S S390C^ in 20 mM Tris pH 7.3, 2 M GuHCl, 1 mM TCEP was mixed with fluorescein maleimide and incubated overnight at room temperature. Free dye was removed using a 3 kDa cutoff Amicon centrifugal concentrator (Merck, NJ), and the buffer was exchanged to 50 mM sodium phosphate, pH 6.5, 150 mM NaCl, 1mM TCEP. Aliquots were stored at -80°C. Fluorescently labeled DNA was purchased from IDT.

### Thioflavin T assays

23 μL samples of ETS DBDs or EWS RNA binding domain constructs (at specified concentrations) and with or without the appropriate DNA oligonucleotide (purchased from IDT, Table 2) were dissolved in 20 mM sodium phosphate buffer pH 7.4, 150 mM NaCl (or 0 mM NaCl), 10 μM ThT and prepared in triplicate in 384-well flat bottom black plates (Greiner Bio-One, NC). Immediately prior to the start of the assay, 2 μL of a 625 μM stock of EWS^LCD^ (or EWS-FLI1 or WT EWS) in 20 mM CAPS pH 11 was added and mixed thoroughly by pipetting to initiate phase separation and the plate was sealed (Thermo Scientific). ThT fluorescence was read at 10-minute intervals for 24 hours at 25°C using a Tecan Infinite M200 plate reader (Tecan Trading AG). Raw fluorescence intensities were normalized to a control sample.

### Pelleting assays

Aged samples (T > 10 hours) from ThT assays were removed from 384-well plates and centrifuged at 21,300 x g for 15 minutes at ambient temperature. The supernatant was removed, and the pellet was resuspended in 8 M urea. The supernatant, pellet, and a sample prior to centrifugation (“total”) were analyzed by SDS-PAGE.

### Microscopy

Freshly prepared (T ~ 0-1 hour) or aged samples (T ~ 24 hours) were imaged directly in sealed 384-well microwell plates with a BioTek Cytation Gen 5 imaging plate reader (Agilent) using a 20 x objective. Alternatively, sealed, chambered (50-well) coverslips (Grace Biolabs), coated with 1% Pluronic F-127 were imaged with an Olympus FV300 inverted confocal microscope using a 40x oil-immersion objective lens operating at 1% power using the 488 nm laser for transmitted light and the 488 nm and 650 nm lasers for fluorescence imaging. Images were acquired simultaneously in differential interference contrast (DIC) and fluorescent modes. Images were processed using Fiji^78^ and fluorescence intensity ratios were extracted by selecting 16 regions outside of the condensed phase and 16 regions inside the condensed phase across 6 – 8 image frames per sample.

To measure fluorescence recovery after photobleaching (FRAP), circular regions of interest were irradiated at a laser power of 10% and fluorescence recovery was monitored at 250-millisecond intervals for 30 seconds after bleaching. Image contrast was adjusted globally. The fluorescence intensities for bleached regions were obtained using Fiji. FRAP curves were normalized using a 0 – 1 scale by setting pre-bleach points to be 1 and the intensity of the first post-bleach point to be 0.

Transmission electron micrographs were acquired on aged samples (T ~ 24 hours) from ThT assay endpoints. Samples were removed from 384-well plates and applied to formvar/carbon coated grids (Electron Microscopy Sciences, PA). Samples were stained with 2% uranyl acetate and imaged using a JEOL 1400 Transmission electron microscope with a XR80 camera (AMT imaging, MA) at a magnification of 50,000 x.

### Nuclear magnetic resonance spectroscopy

NMR experiments were conducted on a Bruker Avance NEO spectrometer (Bruker, MA) operating at a proton Larmor frequency of 700.13 MHz at a temperature of 25°C. ^1^H,^15^N-HSQC, HNCACB, CBCA(CO)NH, HNCO, HN(CA)CO and HCC(CO)NH non-uniformly sampled (15-30% sampling density) data sets were recorded on a of 425 μM sample of ^15^N,^13^C PU. 1 168-270 (Table 2, Supplementary Fig. 1) dissolved in 20 mM sodium phosphate buffer pH 6.4, 0.5 mM EDTA, 0.02% sodium azide. Detailed acquisition parameters are included in the Supplementary Material. Spectra were reconstructed using the SMILE algorithm^79^ and processed with the nmrPipe suite of programs^80^. Backbone ^1^HN, ^13^Ca, ^13^Cβ, ^13^C’ and ^15^N resonances were assigned using CCPNMR Analysis 3.0 software^81^. Acquisition parameters and a detailed description of the measurement of *R1, R2* and heteronuclear NOE is included in the Supplementary Material. Chemical shift perturbations were measured by titrating a sample of 50 μM ^15^N labelled PU.1 168-270 (Table 2, Supplementary Fig. 1) in 50 mM sodium phosphate buffer pH 6.5 with aliquots of a 2 mM stock of EWS^LCD^ in 20 mM CAPS pH 11. The final concentration of the EWS^LCD^ at each titration point was 12.5, 25, 50, 100 and 150 μM resulting in an ~ 7 % dilution of PU.1 across the titration. ^1^H, ^15^N-HSQC spectra were recorded for each titration point processed in Topspin 4.1.1 and analyzed using CCPNMR Analysis 3.0 software. Acquisition parameters and details of CSP calculations are included in the Supplementary Materials.

### Electrophoretic mobility shift assays

20 μL samples were loaded into wells of a 20% polyacrylamide gel prepared in 0.5 x TBE buffer. Electrophoresis was performed for 60 minutes at 150 V using 0.5 x TBE and stained with SYBRsafe (Thermo Fisher, MA, USA) according to the manufacturer’s specifications for 30 minutes before imaging using UV transillumination.

### Circular dichroism

Circular dichroic spectra were recorded on 10 μM samples of all proteins in 50 mM sodium phosphate buffer pH 6.5 at 25°C in a 2 mm pathlength cuvette using a Jasco 810 spectropolarimeter (Jasco, OK) at a scan speed of 50 nm/min with 0.2 nm increments. Each sample was recorded in triplicate, the data was then averaged and converted to mean residue ellipticity.

## Acknowledgements

DSL is the Shohet Family Fund for Ewing Sarcoma Research St. Baldrick’s Scholar and acknowledges the support of the St. Baldrick’s Foundation (634706). EES was supported by a CPRIT Research Training Award (RP170345). This study was funded in part by the Welch Foundation AQ-2001-20190330 (to DSL), NIGMS R01GM140127 (to DSL) and GCCRI Startup Funds (DSL). The authors would like to thank Dr. Dmitri Ivanov for the use of the BioTek Cytation Gen 5 imaging plate reader, Barbara Hunter, Technical Director of the UTHSA Electron Microscopy Laboratory for assistance with transmission electron microscopy, and Dr. Kristin Cano for NMR technical assistance and valuable discussions. This work is based upon research conducted in the Structural Biology Core Facilities, a part of the Institutional Research Cores at the University of Texas Health Science Center at San Antonio supported by the Office of the Vice President for Research and the Mays Cancer Center Drug Discovery and Structural Biology Shared Resource (NIH P30 CA05417X4).

## Data Availability

NMRPipe processing scripts are available upon reasonable request, expression plasmids containing the EWS, EWS-FLI1, and the ETS DBD constructs were deposited with Addgene (#####). The backbone resonance assignments for the PU.1^DBD^ were deposited in the BMRB (#####).

## Author contributions

EES made the protein samples, developed the ThT assays, prepared reagents, collected, processed, and interpreted data, wrote manuscript, conceptualized the study, AKRM made protein samples, conducted phase separation assays, processed data, SA made protein samples, conducted phase separation assays, processed data, XX designed expression construct plasmids, purified protein, DSL collected and processed data, wrote manuscript, obtained funding, and conceptualized the study. All authors contributed to editing the manuscript and have read and approved the manuscript for publication.

## Abbreviations

BAF: ATP-dependent BRG1/BRM associated factor
BRCA1: Breast cancer type 1 susceptibility protein
CSP: chemical shift perturbation
CTD: C-terminal domain
DBD: DNA-binding domain
EDTA: ethylenediaminetetraacetic acid
ERG: Transcriptional regulator ERG
ETV1: ETS translocation variant 1
ETS: E-twenty-six transformation-specific
EWS: RNA-binding protein EWS
EwS: Ewing sarcoma
FEV: protein FEV
FLI1: Friend leukemia integration 1
FRAP: fluorescence recovery after photobleaching
FUS: fused in sarcoma
GFP: green fluorescent protein
HA: high-affinity
HSQC: heteronuclear single quantum coherence
IDR: intrinsically disordered region
LCD: low-complexity domain
LLPS: liquid-liquid phase separation
NMR: nuclear magnetic resonance
PNET: primitive neuroectodermal tumor
PU.1: transcription factor PU.1
RNA Pol II: DNA-directed RNA polymerase II subunit RPB1
RGG: Arg-Gly-Gly
RRM: RNA-recognition motif
SEC: size-exclusion chromatography
TAF15: TATA-binding protein associated factor 2N
TEM: transmission electron microscopy
TEV: Tobacco Etch Virus
ThT: Thioflavin T
WT: wild-type

## References

1. Grunewald, T.G.P. et al. Ewing sarcoma. Nat Rev Dis Primers 4, 5 (2018).

2. Gorthi, A. et al. EWS-FLI1 increases transcription to cause R-loops and block BRCA1 repair in Ewing sarcoma. Nature 555, 387–391 (2018).

3. Yang, L., Chansky, H.A. & Hickstein, D.D. EWS.Fli-1 fusion protein interacts with hyperphosphorylated RNA polymerase II and interferes with serine-arginine protein-mediated RNA splicing. J Biol Chem 275, 37612–8 (2000).

4. Boone, M.A. et al. The FLI portion of EWS/FLI contributes a transcriptional regulatory function that is distinct and separable from its DNA-binding function in Ewing sarcoma. Oncogene 40, 4759–4769 (2021).

5. Jaishankar, S., Zhang, J., Roussel, M.F. & Baker, S.J. Transforming activity of EWS/FLI is not strictly dependent upon DNA-binding activity. Oncogene 18, 5592–7 (1999).

6. Kedage, V. et al. An Interaction with Ewing’s Sarcoma Breakpoint Protein EWS Defines a Specific Oncogenic Mechanism of ETS Factors Rearranged in Prostate Cancer. Cell Rep 17, 1289–1301 (2016).

7. Panagopoulos, I. et al. Fusion of the FUS gene with ERG in acute myeloid leukemia with t(16;21)(p11;q22). Genes Chromosomes Cancer 11, 256–62 (1994).

8. Aman, P. et al. Rearrangement of the transcription factor gene CHOP in myxoid liposarcomas with t(12;16)(q13;p11). Genes Chromosomes Cancer 5, 278–85 (1992).

9. Martini, A. et al. Recurrent rearrangement of the Ewing’s sarcoma gene, EWSR1, or its homologue, TAF15, with the transcription factor CIZ/NMP4 in acute leukemia. Cancer Res 62, 5408–12 (2002).

10. Kato, M. et al. Cell-free formation of RNA granules: low complexity sequence domains form dynamic fibers within hydrogels. Cell 149, 753–67 (2012).

11. Kwon, I. et al. Phosphorylation-regulated binding of RNA polymerase II to fibrous polymers of low-complexity domains. Cell 155, 1049–1060 (2013).

12. Chong, S. et al. Imaging dynamic and selective low-complexity domain interactions that control gene transcription. Science 361(2018).

13. Hollenhorst, P.C., McIntosh, L.P. & Graves, B.J. Genomic and biochemical insights into the specificity of ETS transcription factors. Annu Rev Biochem 80, 437–71 (2011).

14. Oh, S., Shin, S. & Janknecht, R. ETV1, 4 and 5: an oncogenic subfamily of ETS transcription factors. Biochim Biophys Acta 1826, 1–12 (2012).

15. Sizemore, G.M., Pitarresi, J.R., Balakrishnan, S. & Ostrowski, M.C. The ETS family of oncogenic transcription factors in solid tumours. Nat Rev Cancer 17, 337–351 (2017).

16. Gutierrez-Hartmann, A., Duval, D.L. & Bradford, A.P. ETS transcription factors in endocrine systems. Trends Endocrinol Metab 18, 150–8 (2007).

17. Hsing, M., Wang, Y., Rennie, P.S., Cox, M.E. & Cherkasov, A. ETS transcription factors as emerging drug targets in cancer. Med Res Rev 40, 413–430 (2020).

18. Fisher, C. The diversity of soft tissue tumours with EWSR1 gene rearrangements: a review. Histopathology 64, 134–50 (2014).

19. Boulay, G. et al. Cancer-Specific Retargeting of BAF Complexes by a Prion-like Domain. Cell 171, 163–178 e19 (2017).

20. Zuo, L. et al. Loci-specific phase separation of FET fusion oncoproteins promotes gene transcription. Nat Commun 12, 1491 (2021).

21. Petermann, R. et al. Oncogenic EWS-Fli1 interacts with hsRPB7, a subunit of human RNA polymerase II. Oncogene 17, 603–10 (1998).

22. May, W.A. et al. Ewing sarcoma 11;22 translocation produces a chimeric transcription factor that requires the DNA-binding domain encoded by FLI1 for transformation. Proc Natl Acad Sci U S A 90, 5752–6 (1993).

23. May, W.A. et al. The Ewing’s sarcoma EWS/FLI-1 fusion gene encodes a more potent transcriptional activator and is a more powerful transforming gene than FLI-1. Mol Cell Biol 13, 7393–8 (1993).

24. Johnson, K.M. et al. Role for the EWS domain of EWS/FLI in binding GGAA-microsatellites required for Ewing sarcoma anchorage independent growth. Proc Natl Acad Sci U S A 114, 9870–9875 (2017).

25. Lessnick, S.L., Braun, B.S., Denny, C.T. & May, W.A. Multiple domains mediate transformation by the Ewing’s sarcoma EWS/FLI-1 fusion gene. Oncogene 10, 423–31 (1995).

26. Gangwal, K., Close, D., Enriquez, C.A., Hill, C.P. & Lessnick, S.L. Emergent Properties of EWS/FLI Regulation via GGAA Microsatellites in Ewing’s Sarcoma. Genes Cancer 1, 177–187 (2010).

27. Gangwal, K. et al. Microsatellites as EWS/FLI response elements in Ewing’s sarcoma. Proc Natl Acad Sci U S A 105, 10149–54 (2008).

28. Weber, S.C. & Brangwynne, C.P. Getting RNA and protein in phase. Cell 149, 1188–91 (2012).

29. Boeynaems, S. et al. Protein Phase Separation: A New Phase in Cell Biology. Trends Cell Biol (2018).

30. Mitrea, D.M. & Kriwacki, R.W. Phase separation in biology; functional organization of a higher order. Cell Commun Signal 14, 1 (2016).

31. Hnisz, D., Shrinivas, K., Young, R.A., Chakraborty, A.K. & Sharp, P.A. A Phase Separation Model for Transcriptional Control. Cell 169, 13–23 (2017).

32. Boehning, M. et al. RNA polymerase II clustering through carboxy-terminal domain phase separation. Nat Struct Mol Biol 25, 833–840 (2018).

33. Liao, S.E. & Regev, O. Splicing at the phase-separated nuclear speckle interface: a model. Nucleic Acids Res 49, 636–645 (2021).

34. Strom, A.R. et al. Phase separation drives heterochromatin domain formation. Nature 547, 241–245 (2017).

35. Protter, D.S. & Parker, R. Principles and Properties of Stress Granules. Trends Cell Biol 26, 668–79 (2016).

36. Spahn, L. et al. Homotypic and heterotypic interactions of EWS, FLI1 and their oncogenic fusion protein. Oncogene 22, 6819–29 (2003).

37. Ahmed, N.S. et al. Fusion protein EWS-FLI1 is incorporated into a protein granule in cells. RNA (2021).

38. Murthy, A.C. et al. Molecular interactions contributing to FUS SYGQ LC-RGG phase separation and co-partitioning with RNA polymerase II heptads. Nat Struct Mol Biol 28, 923–935 (2021).

39. Qamar, S. et al. FUS Phase Separation Is Modulated by a Molecular Chaperone and Methylation of Arginine Cation-pi Interactions. Cell 173, 720–734 e15 (2018).

40. Chong, S. et al. Tuning levels of low-complexity domain interactions to modulate endogenous oncogenic transcription. Mol Cell (2022).

41. Patel, A. et al. A Liquid-to-Solid Phase Transition of the ALS Protein FUS Accelerated by Disease Mutation. Cell 162, 1066–77 (2015).

42. Murray, D.T. et al. Structure of FUS Protein Fibrils and Its Relevance to Self-Assembly and Phase Separation of Low-Complexity Domains. Cell 171, 615–627 e16 (2017).

43. Kato, M. & McKnight, S.L. The low-complexity domain of the FUS RNA binding protein self-assembles via the mutually exclusive use of two distinct cross-beta cores. Proc Natl Acad Sci U S A 118(2021).

44. Liu, Z. et al. Hsp27 chaperones FUS phase separation under the modulation of stress-induced phosphorylation. Nat Struct Mol Biol 27, 363–372 (2020).

45. Guo, L. et al. Nuclear-Import Receptors Reverse Aberrant Phase Transitions of RNA-Binding Proteins with Prion-like Domains. Cell 173, 677–692 e20 (2018).

46. Couthouis, J. et al. Evaluating the role of the FUS/TLS-related gene EWSR1 in amyotrophic lateral sclerosis. Hum Mol Genet 21, 2899–911 (2012).

47. Karim, F.D. et al. The ETS-domain: a new DNA-binding motif that recognizes a purine-rich core DNA sequence. Genes Dev 4, 1451–3 (1990).

48. Guillon, N. et al. The oncogenic EWS-FLI1 protein binds in vivo GGAA microsatellite sequences with potential transcriptional activation function. PLoS One 4, e4932 (2009).

49. Hou, C., Mandal, A., Rohr, J. & Tsodikov, O.V. Allosteric interference in oncogenic FLI1 and ERG transactions by mithramycins. Structure 29, 404–412 e4 (2021).

50. Hou, C. & Tsodikov, O.V. Structural Basis for Dimerization and DNA Binding of Transcription Factor FLI1. Biochemistry 54, 7365–74 (2015).

51. Kodandapani, R. et al. A new pattern for helix-turn-helix recognition revealed by the PU.1 ETS-domain-DNA complex. Nature 380, 456–60 (1996).

52. Regan, M.C. et al. Structural and dynamic studies of the transcription factor ERG reveal DNA binding is allosterically autoinhibited. Proc Natl Acad Sci U S A 110, 13374–9 (2013).

53. Cooper, C.D., Newman, J.A., Aitkenhead, H., Allerston, C.K. & Gileadi, O. Structures of the Ets Protein DNA-binding Domains of Transcription Factors Etv1, Etv4, Etv5, and Fev: DETERMINANTS OF DNA BINDING AND REDOX REGULATION BY DISULFIDE BOND FORMATION. J Biol Chem 290, 13692–709 (2015).

54. Liang, H. et al. The secondary structure of the ets domain of human Fli-1 resembles that of the helix-turn-helix DNA-binding motif of the Escherichia coli catabolite gene activator protein. Proc Natl Acad Sci U S A 91, 11655–9 (1994).

55. Laudet, V., Hanni, C., Stehelin, D. & Duterque-Coquillaud, M. Molecular phylogeny of the ETS gene family. Oncogene 18, 1351–9 (1999).

56. Mao, X., Miesfeldt, S., Yang, H., Leiden, J.M. & Thompson, C.B. The FLI-1 and chimeric EWS-FLI-1 oncoproteins display similar DNA binding specificities. J Biol Chem 269, 18216–22 (1994).

57. Johnson, C.N., Xu, X., Holloway, S.P. & Libich, D.S. The (1)H, (15)N and (13)C resonance assignments of the low-complexity domain from the oncogenic fusion protein EWS-FLI1. Biomol NMR Assign 16, 67–73 (2022).

58. Jumper, J. et al. Highly accurate protein structure prediction with AlphaFold. Nature 596, 583–589 (2021).

59. Li, H. et al. Ewing sarcoma gene EWS is essential for meiosis and B lymphocyte development. J Clin Invest 117, 1314–23 (2007).

60. Paronetto, M.P., Minana, B. & Valcarcel, J. The Ewing sarcoma protein regulates DNA damage-induced alternative splicing. Mol Cell 43, 353–68 (2011).

61. Knoop, L.L. & Baker, S.J. The splicing factor U1C represses EWS/FLI-mediated transactivation. J Biol Chem 275, 24865–71 (2000).

62. Embree, L.J., Azuma, M. & Hickstein, D.D. Ewing sarcoma fusion protein EWSR1/FLI1 interacts with EWSR1 leading to mitotic defects in zebrafish embryos and human cell lines. Cancer Res 69, 4363–71 (2009).

63. Hallier, M., Lerga, A., Barnache, S., Tavitian, A. & Moreau-Gachelin, F. The transcription factor Spi-1/PU.1 interacts with the potential splicing factor TLS. J Biol Chem 273, 4838–42 (1998).

64. Delva, L. et al. Multiple functional domains of the oncoproteins Spi-1/PU.1 and TLS are involved in their opposite splicing effects in erythroleukemic cells. Oncogene 23, 4389–99 (2004).

65. Janke, A.M. et al. Lysines in the RNA Polymerase II C-Terminal Domain Contribute to TAF15 Fibril Recruitment. Biochemistry (2017).

66. Kato, M. & McKnight, S.L. Cross-beta Polymerization of Low Complexity Sequence Domains. Cold Spring Harb Perspect Biol 9(2017).

67. Hughes, M.P. et al. Atomic structures of low-complexity protein segments reveal kinked beta sheets that assemble networks. Science 359, 698–701 (2018).

68. Lee, M., Ghosh, U., Thurber, K.R., Kato, M. & Tycko, R. Molecular structure and interactions within amyloid-like fibrils formed by a low-complexity protein sequence from FUS. Nat Commun 11, 5735 (2020).

69. Sawaya, M.R., Hughes, M.P., Rodriguez, J.A., Riek, R. & Eisenberg, D.S. The expanding amyloid family: Structure, stability, function, and pathogenesis. Cell 184, 4857–4873 (2021).

70. Ng, K.P. et al. Multiple aromatic side chains within a disordered structure are critical for transcription and transforming activity of EWS family oncoproteins. Proc Natl Acad Sci U S A 104, 479–84 (2007).

71. Murthy, A.C. et al. Molecular interactions underlying liquid-liquid phase separation of the FUS low-complexity domain. Nat Struct Mol Biol 26, 637–648 (2019).

72. Ryan, J.J., Sprunger, M.L., Holthaus, K., Shorter, J. & Jackrel, M.E. Engineered protein disaggregases mitigate toxicity of aberrant prion-like fusion proteins underlying sarcoma. J Biol Chem 294, 11286–11296 (2019).

73. Selvanathan, S.P. et al. Oncogenic fusion protein EWS-FLI1 is a network hub that regulates alternative splicing. Proc Natl Acad Sci U S A 112, E1307–16 (2015).

74. Bertolotti, A. et al. EWS, but not EWS-FLI-1, is associated with both TFIID and RNA polymerase II: interactions between two members of the TET family, EWS and hTAFII68, and subunits of TFIID and RNA polymerase II complexes. Mol Cell Biol 18, 1489–97 (1998).

75. Kapust, R.B. et al. Tobacco etch virus protease: mechanism of autolysis and rational design of stable mutants with wild-type catalytic proficiency. Protein Eng 14, 993–1000 (2001).

76. Burke, K.A., Janke, A.M., Rhine, C.L. & Fawzi, N.L. Residue-by-Residue View of In Vitro FUS Granules that Bind the C-Terminal Domain of RNA Polymerase II. Mol Cell 60, 231–41 (2015).

77. Mao, H., Hart, S.A., Schink, A. & Pollok, B.A. Sortase-mediated protein ligation: a new method for protein engineering. J Am Chem Soc 126, 2670–1 (2004).

78. Schindelin, J. et al. Fiji: an open-source platform for biological-image analysis. Nat Methods 9, 676–82 (2012).

79. Ying, J., Delaglio, F., Torchia, D.A. & Bax, A. Sparse multidimensional iterative lineshape-enhanced (SMILE) reconstruction of both non-uniformly sampled and conventional NMR data. J Biomol NMR 68, 101–118 (2017).

80. Delaglio, F. et al. NMRPipe: a multidimensional spectral processing system based on UNIX pipes. J Biomol NMR 6, 277–93 (1995).

81. Skinner, S.P. et al. CcpNmr AnalysisAssign: a flexible platform for integrated NMR analysis. J Biomol NMR 66, 111–124 (2016).

