## Supplemental Materials for "The oncogenic fusion protein EWS-FLI1 promotes premature ageing of biomolecular condensates by catalyzing fibril formation"

**Supplementary Materials and Methods**

**Recombinant Protein Cloning.** All gene constructs were codon optimized for *E. coli* expression, synthesized (GenScript) and cloned into modified pET expression vectors with an 8 x His-tag followed by a tobacco etch virus (TEV) protease cleavage site, both N-terminal to the protein coding sequence except for full-length FUS which was cloned into a pTHMT vector with an N-terminal MBP tag[1]. The F385A mutation in EWS-FLI1 and the F362A mutation in $FLI1^{DBD}$ correspond to the same residue in FLI1 but with numbering corresponding to EWS-FLI numbering or FLI1 numbering, respectively. This mutation was used for this study because it was shown to reduce the propensity of $FLI1^{DBD}$ to dimerize[2]. All expression plasmids were verified by DNA sequencing. Full-length MBP-FUS was a gift from Nicolas Fawzi (Addgene plasmid #98651; http://n2t.net/addgene:98651 ; RRID:Addgene_98651)

**Recombinant Protein Expression.** Plasmids were transformed using the heat-shock method. *E. coli* BL21 Star™ (DE3) (Invitrogen, MA) cells were used for expression of the following proteins: full-length FUS, full-length EWS, full-length EWS-FLI1, $EWS^{LCD}$, EWS-FLI1 194-445 F385A, FLI1 276-399 F362A ($FLI1^{DBD}$), FLI1 276-361 ($FLI1^{DBD\ \Delta\alpha4}$), $FLI1^{DBD\ C299S\ S390C}$, ERG 306-429 F392A ($ERG^{DBD}$), EWS 359-447 ($EWS^{RRM}$) and EWS 359-513 ($EWS^{RRM\text{-}RGG2}$) (Table 1, Supplementary Fig. 1). BL21 (DE3) pLysS competent cells (Invitrogen) were used for expression of the following proteins: $FLI1^{DBD}$ $^{R337A\ R340A}$ ($FLI1^{DBD\ R2L2}$), ETV1 332-458 ($ETV1^{DBD}$) and PU.1 168-270 ($PU.1^{DBD}$). To produce proteins without isotopic enrichment, one colony of the resulting plate was used to inoculate a 100 mL LB starter culture that was grown at 37°C overnight with shaking at 225 rpm. For expression of isotopically enriched proteins, one colony was used to inoculate a 2 mL LB starter culture that was grown at 37 °C with 225 rpm shaking for ~ 5-6 hours. 1 mL of this culture was used to inoculate a 100 mL M9 starter culture which was grown overnight at 37°C with shaking at 225 rpm. M9 media was supplemented with $^{15}NH_4Cl$ (1g/L) and 0.02% (w/v) yeast extract for $^{15}N$ labelling or $^{15}NH_4Cl$ and $^{13}C_6$ D-glucose (3 g/L) with 0.02% (w/v) Isogro®-$^{13}C$, $^{15}N$ (Sigma, MO) for $^{15}N$, $^{13}C$ labelling. All cultures were also supplemented with 100 μg/mL ampicillin or 50 μg/mL kanamycin for FUS constructs. 25 – 50 mL of the overnight cultures were used to inoculate 1 L LB or M9 cultures, which were grown in baffled Fernbach flasks at 37 °C with shaking at 225 rpm until $OD_{600}$ reached ~ 0.6-0.8, protein expression was induced with 1 mM IPTG. Protein expression was continued for 3 hours at 37 °C for EWS 359-447, or for 4 hours at 37 °C for full-length FUS, full-length EWS, full-length EWS-FLI1, EWS-FLI1 194-445 F385A and $ETV1^{DBD}$. For $EWS^{RRM\text{-}RGG2}$, protein expression was continued for 1 hour at 30 °C and for a further 16 hours at 12 °C. For $FLI1^{DBD}$, $FLI1^{DBD\ \Delta\alpha4}$, $FLI1^{DBD\ C299S\ S390C}$, $ERG^{DBD}$, $FLI1^{DBD\ R2L2}$ and $PU.1^{DBD}$, protein expression was continued at 12 °C for 20-24 hours. Subsequently, the cells were harvested by centrifugation at 4000 x g for 30 minutes at 4 °C. Pellets were either frozen directly at -20 °C or resuspended in the suitable buffer for purification before being frozen at -20 °C as will be described below.

**Recombinant Protein Purification.** Full-length MBP-FUS was purified as described previously[1]. $EWS^{LCD}$, EWS and EWS-FLI1 were purified as described for $EWS^{LCD}$ in our previous work[3].

EWS-FLI1 194-445 F385A pellets were thawed and resuspended in lysis buffer (20 mM Tris pH 8, 2 M GuHCl, 2 mM DTT). The resuspension was sonicated using a 550 Sonic Dismembrator (Fisher Scientific, PA) for 5 seconds on pulses and 55 second off pulses on ice for a total of 6 min processing time at a power setting of 8 using a ½ inch probe. The suspension was then clarified by centrifugation at 45,000 x g for 30 min at 4 °C. The lysate was filtered through a 0.45 μm syringe-driven filter then applied to applied to a 5 mL HisTrap HP column (Cytiva, MA) and the protein was eluted using a linear gradient from 0 – 500 mM imidazole over 125 mL. Fractions containing EWS-FLI1 194-445 F385A were concentrated to ~ 5 mL and loaded onto a HiLoad 16/600 Superdex 75 pg column (Cytiva, MA) equilibrated with 20 mM Tris pH 7, 2 M GuHCl, 2 mM TCEP. Fractions containing EWS-FLI1 194-445 F385A were pooled and concentrated to 1.2 mM, aliquoted and stored at -80 °C.

Cells expressing FLI1$^{DBD}$, FLI1$^{DBD\ \Delta\alpha4}$ and ERG$^{DBD}$ were resuspended in 50 mM sodium phosphate buffer pH 6.5, 2 mM DTT and frozen at -20 °C until purification. The cells were thawed and sonicated for a total processing time of 3 min as described above and the lysate was further clarified by centrifugation at 45,000 x g for 30 min at 4°C. The supernatant was applied to 5 mL HiTrap SP Sepharose FF column (Cytiva, MA). The column was subsequently washed with ~ 10 column volumes (CV) buffer A (50 mM sodium phosphate buffer pH 6.5, 2 mM DTT). A gradient of 0-30% buffer B (50 mM sodium phosphate buffer pH 6.5, 2 mM DTT, 1 M NaCl) over 10 CVs, 30-55% buffer B over 10 CVs and 55-100% buffer B over 6 CVs was used for elution. Fractions containing the protein of interest were pooled and concentrated at 4 °C with a 3 kDa cutoff Amicon centrifugal concentrator (Merck, NJ) and applied to a HiLoad 16/600 Superdex 75 pg column (Cytiva, MA) equilibrated in 50 mM sodium phosphate buffer, 150 mM NaCl, 1 mM EDTA, 1 mM PMSF, 2 mM TCEP. Fractions containing the protein of interest were pooled, concentrated to 100-300 µM, aliquoted and stored at -80 °C.

Cells expressing FLI1$^{DBD\ C299S\ S390C}$ were frozen as a pellet at – 20 °C until use. Cells were thawed and resuspended in 50 mM Tris pH 8, 500 mM NaCl, 20 mM imidazole, 2 M GuHCl, 2 mM DTT (buffer A). The resuspension was sonicated for a total processing time of 3 mins as described above and clarified by centrifugation at 45,000 x g for 30 min at 4°C. The supernatant was applied to 5 mL HisTrap HP column (Cytiva, MA). The column was washed with 50 mL buffer A and 30 mL of 50 mM Tris pH 8, 500 mM NaCl, 40 mM imidazole, 2 M GuHCl, 2 mM DTT. FLI1$^{DBD\ C299S\ S390C}$ was eluted with 50 mM Tris pH 8, 500 mM NaCl, 500 mM imidazole, 2 M GuHCl, 2 mM DTT. Eluted protein was concentrated to ~ 2 mL with a 3 kDa cutoff Amicon centrifugal concentrator (Merck, NJ) and loaded onto a HiLoad 16/600 Superdex 75 pg column (Cytiva, MA) equilibrated in 20 mM Tris pH 7.3, 2 M GuHCl, 1 mM TCEP. Fractions containing FLI1$^{DBD\ C299S\ S390C}$ were pooled and concentrated to ~80 µM and stored at 4 °C.

Cells expressing FLI1$^{DBD\ R2L2}$ were resuspended in 50 mM Tris pH 8.8, 10 mM imidazole, 300 mM NaCl, 2 mM DTT and frozen at -20°C until use. Cells were thawed and the suspension was sonicated for a total processing time of 3 min as described above and clarified by centrifugation at 45,000 x g for 30 min at 4 °C. The supernatant was loaded onto a 5 mL HisTrap column (Cytiva, MA) and the column was washed with 25 mL of each of: 10 mM imidazole, 20 mM imidazole, 50 mM imidazole prepared in the above buffer. The sample was eluted with 15 mL of 50 mM Tris pH 7.5, 300 mM NaCl, 300 mM imidazole, 2 mM DTT. The eluate was concentrated using a 3 kDa cutoff Amicon centrifugal concentrator (Merck, NJ) and loaded onto a HiLoad 16/600 Superdex 75 pg column (Cytiva, MA) equilibrated with 50 mM sodium phosphate buffer pH 6, 150 mM NaCl, 2 mM TCEP, 0.2 mM PMSF, 0.5 mM EDTA. Fractions containing FLI1$^{DBD\ R2L2}$ were pooled and concentrated to ~ 60 µM, aggregates were removed by centrifugation, aliquoted and stored at -80 °C.

Cells expressing ETV1$^{DBD}$ were resuspended in 50 mM Tris pH 8, 500 mM NaCl, 20 mM imidazole, 2 mM TCEP and stored at -20 °C until use. Cells were thawed and the suspension was sonicated for a total processing time of 3 minutes as described above and the lysate was further clarified by centrifugation at 45,000 x g for 30 min at 4 °C. The supernatant was applied to 5 mL HisTrap HP column (Cytiva, MA) which was then washed with 30 mL of 40 mM imidazole prepared in the above buffer and ETV1$^{DBD}$ was eluted with 500 mM imidazole prepared in the above buffer. The eluted protein was concentrated and loaded onto a HiLoad 16/600 Superdex 75 pg column (Cytiva) equilibrated with 50 mM sodium phosphate buffer pH 7, 150 mM NaCl, 0.5 mM EDTA, 0.2 mM PMSF, 2 mM TCEP. Fractions containing ETV1$^{DBD}$ were pooled and concentrated to ~ 500 μM, aliquoted and stored at -80 °C.

Cells expressing PU.1$^{DBD}$ were resuspended in 50 mM Tris pH 8, 500 mM NaCl, 20 mM imidazole and stored at -20 °C until use. Cells were thawed and sonicated for a total processing time of 3 minutes and the lysate was further clarified by centrifugation at 45, 000 x g for 30 min at 4 °C. The supernatant was loaded onto a 5 mL HisTrap HP column (Cytiva, MA). The column was washed with 50 mL 20 mM imidazole prepared in the above buffer, 50 mL 20 mM imidazole + 1 M NaCl prepared in the above buffer, 30 mL 50 mM imidazole prepared in the above buffer and PU.1$^{DBD}$ was eluted with 500 mM imidazole prepared in the above buffer. To confirm that the effects of ETS proteins on EWS$^{LCD}$ condensates were not due to the 8 x His-tag that was not cleaved from FLI1$^{DBD}$, ERG$^{DBD}$ or ETV1$^{DBD}$, PU.1$^{DBD}$ was purified both with an 8 x His-tag and with the tag cleaved. For the purification of His-tagged PU.1$^{DBD}$, eluted protein was concentrated to ~ 10 mL and loaded onto a HiLoad 16/600 Superdex 75 pg column (Cytiva, MA) equilibrated in 50 mM sodium phosphate buffer pH 6.5, 150 mM NaCl, 0.5 mM EDTA, 0.2 mM PMSF, 2 mM TCEP. Fractions containing PU.1$^{DBD}$ were pooled and concentrated to ~ 150 μM and stored as aliquots at -80 °C. For the purification of PU.1$^{DBD}$ with tag cleavage, the HisTrap elution was diluted six-fold in 50 mM Tris pH 8 (heparin buffer A) and loaded onto a 5 mL HiTrap Heparin HP column (Cytiva, MA). The column was washed with heparin buffer A until the UV absorbance stopped decreasing. PU.1$^{DBD}$ was eluted from the heparin column with heparin buffer A + 2 M NaCl. Eluted protein was dialyzed against 1 L 50 mM Tris pH 8, 1 mM DTT, 0.5 mM EDTA, 100 mM NaCl, 1 M urea for 4 hours at room temperature. 1 mL TEV was added (A280 = 1.25) and the sample was dialyzed against a fresh 1 L of the same buffer at room temperature overnight. Aggregates were removed by centrifugation and the sample was concentrated and applied to a HiLoad 16/600 Superdex 75 pg column (Cytiva, MA) equilibrated with 50 mM Tris pH 8, 500 mM NaCl. Fractions containing cleaved PU.1$^{DBD}$ were pooled and dialyzed against 2 L 50 mM sodium phosphate buffer pH 6.5 for ~ 4 hours at 4 °C. The dialysate was concentrated to ~90 μM, aliquoted and stored at -80 °C.

Cells expressing EWS$^{RRM}$ and EWS$^{RRM\text{-}RGG2}$ were resuspended in 50 mM Tris pH 8, 500 mM NaCl, 20 mM imidazole, 5 mM DTT. The suspension was sonicated for a total processing time of 3 minutes as described above and further clarified by centrifugation at 45,000 x g for 30 min at 4 °C. The supernatant was applied to a 5 mL HisTrap HP column (Cytiva, MA). The column was washed with 50 mL 20 mM imidazole, 30 mL 40 mM imidazole, 30 mL 50 mM imidazole, all prepared in the above buffer and EWS$^{RRM}$/EWS$^{RRM\text{-}RGG2}$ were eluted with 500 mM imidazole prepared in the above buffer. The elution was concentrated and applied to a HiLoad 16/600 Superdex 75 pg column (Cytiva, MA) equilibrated with 20 mM Tris pH 7.0, 100 mM NaCl (for EWS$^{RRM}$) or 50 mM NaCl (for EWS$^{RRM\text{-}RGG2}$), 2 mM TCEP, 0.02% sodium azide. Fractions containing EWS$^{RRM}$ or EWS$^{RRM\text{-}RGG2}$ were pooled and concentrated to ~ 700-800 μM, aliquoted and stored at -80 °C.

**Fluorescent labelling using sortase.** Purified EWS$^{LCD}$, EWS and EWS-FLI1 were fluorescently labelled with DyLight 488, Dylight 650 or fluorescein maleimide 488 (Thermo Scientific, MA) using sortase[4] and the SortC1 peptide, KLPETGG (GenScript, NJ). The dyes were resolubilized according to the manufacturer's specifications in DMF to 1 mg/mL and 10 mg/mL for the 488 and 650 dyes, respectively.

Solutions containing the dyes were then mixed with a 3 to 5-fold molar excess of the SortC1 peptide dissolved in 0.1 M sodium bicarbonate buffer at pH 9.5 at room temperature for one hour. Aliquots were frozen at -80 °C until use. 50 - 100 μM solutions of the proteins to be fluorescently labelled were prepared in 0.1 M sodium bicarbonate pH 9.5 with 50-100 μM of the dye-labelled peptides and 2.5 μM of the sortase enzyme. These reactions were incubated overnight at room temperature protected from light. Fluorescent labelling of the protein of interest was verified by SDS-PAGE followed by UV transillumination, and the free dye, peptide and enzyme was removed by SEC using a HiLoad 16/600 Superdex 75 pg column (Cytiva, MA). Fractions containing the fluorescently labelled protein were pooled, concentrated and stored at -80 °C until use.

**Fluorescein maleimide labelling of FLI1$^{DBD\ C299S\ S390C}$.** 2 mg of fluorescein maleimide dye was added to ~ 4 mL of ~ 80 μM FLI1$^{DBD\ C299S\ S390C}$ in 20 mM Tris pH 7.3, 2 M GuHCl, 1 mM TCEP and this mixture was incubated overnight at room temperature with gentle shaking. Free dye was removed via buffer exchange using a 3 kDa cutoff Amicon centrifugal concentrator (Merck, NJ) and a final buffer exchange step into 50 mM sodium phosphate pH 6.5 150 mM NaCl, 1mM TCEP was used to remove GuHCl and induce refolding. Aggregates were removed via centrifugation and aliquots were stored at -80 °C at ~ 30 μM.

**Thioflavin T assays.** For all ThT assays, 25 μL samples were prepared in triplicate in 384-well flat bottom black plates (Greiner Bio-One, NC). 2.5 μL of 100 μM ThT prepared in 50 mM Tris pH 7.5 was added to each well to achieve a final ThT concentration of 10 μM. 1.9 μL of 2 M NaCl was added to each well to achieve a final NaCl concentration of 150 mM unless otherwise specified. For most assays, 50 μM stocks of ETS proteins and control proteins and 50 μM stocks of DNA (IDT, IA; Table 2) were prepared in the assay buffer (20 mM sodium phosphate buffer pH 7.4 + 0.02% sodium azide) and 2.5 μL of these stocks were added to the indicated samples. For assays in which higher concentrations of ETS proteins, control proteins or DNA was required, more concentrated stocks were used such that the total volume of the protein/DNA stock required per sample was < 7 μL. To ensure that the observed effects were not due to the different storage buffers used for the various ETS proteins and control proteins, ThT assays have been conducted using Tris, MES and sodium phosphate buffers, yielding similar results (data not shown). Since the majority of the ETS proteins were stored in sodium phosphate buffers, the volume of each sample was adjusted to 23 μL using the assay buffer specified above and then mixed thoroughly. Immediately prior to the start of the assay, 2 μL of a 625 μM stock of EWS$^{LCD}$ in 20 mM CAPS pH 11 was added to each sample to initiate phase separation and the sample was mixed thoroughly by pipetting. The plate was sealed using clear sealing tape (Thermo Scientific, MA). ThT fluorescence was read at 3-minute intervals with 10 seconds of shaking prior to each read over 24 hours at 25 °C using 440 nm excitation and 485 nm emission wavelengths on a Tecan infinite M200 plate reader (Tecan Trading AG). Raw fluorescence intensities for each experiment were normalized to a 0 – 1.0 scale by setting the maximum ThT fluorescence value obtained for 50 μM EWS$^{LCD}$ + 5 μM FLI1$^{DBD}$ after 10 hours to be 1 and the lowest ThT fluorescence observed for any sample across the assay at 0 hours to be 0.

**Pelleting assays.** Aged samples (T > 10 hours) from ThT assays were removed from 384-well plates and centrifuged at 21,300 x g for 15 minutes at ambient temperature. The supernatant was removed, and the pellet was resuspended in 8 M urea. The supernatant, pellet, and a sample prior to centrifugation ("total") were analyzed by SDS-PAGE.

**Bright-field microscopy.** Freshly prepared (T ~0-1 hour) or aged samples (T ~ 24 hours) were added to wells of a 384-well flat bottom black plate (Greiner Bio-One, NC) and the plate was sealed with clear sealing tape (Thermo Scientific, MA). Wells were imaged on a BioTek Cytation Gen 5 imaging plate reader (Agilent, CA) using an objective magnification of 20 x.

**DIC and fluorescence microscopy.** 25 μL samples were prepared with various mixtures of $EWS^{LCD}$, $FLI1^{DBD}$ and HA DNA. In the indicated samples, 650-labelled EWS, 488-labelled EWS, 650-labelled HA DNA and 488-labelled $FLI1^{DBD}$ were used at a final concentration of 0.5 μM. Appropriate concentrations of stocks of unlabeled $EWS^{LCD}$, $FLI1^{DBD}$ and HA DNA were used to reach the required final concentration. Because only two fluorescent labels were used, samples were prepared with two labelled components and one unlabeled component. The unlabeled component was either $FLI1^{DBD}$ or HA DNA, while $EWS^{LCD}$ was always fluorescently labeled with either the 488 or 650 dye. As for the ThT assays, 2 M NaCl was added to a final concentration of 150 mM. Appropriate volumes of ETS proteins/control proteins/DNA stocks and appropriate volumes of assay buffer (see above) were added to adjust the sample volume to 23 μL and $EWS^{LCD}$ was added as the final component after thorough mixing. 4.5 μL samples were applied to freshly coated (1% Pluronic F-127), chambered (50-well) coverslips (Grace Biolabs, OR). Chambers were sealed with a second coverslip to reduce evaporation and incubated for 10 min at ambient temperature before imaging using an Olympus FV300 inverted confocal microscope (Olympus). A 40 x oil-immersion objective lens operating at 1% power using the 488 nm laser was used for transmitted light and the 488 nm and 650 nm lasers for fluorescence imaging. Images were acquired simultaneously in differential interference contrast (DIC) and fluorescent modes.

**Calculation of fluorescence intensity ratios.** 25 μL samples were prepared with 0.5 μM $FLI1^{DBD\ 488}$ and $FLI1^{DBD}$ to a final concentration of 5 μM. Various amounts of unlabeled HA DNA were added to achieve the desired final DNA concentration. $EWS^{LCD\ 650}$ was added to a final concentration of 0.5 μM and a 625 μM stock of unlabeled EWS was added as the final component after thorough mixing to achieve a total $EWS^{LCD}$ concentration of 50 μM. Samples were added to 384-well flat bottom black plates (Greiner Bio-One, NC) and the plates were sealed with clear sealing tape (Thermo Scientific, MA). Samples were imaged as described above. Fluorescence intensity ratios were obtained by selecting 16 regions outside of the condensed phase and 16 regions inside the condensed phase across 6 – 8 image frames taken of each sample. The raw fluorescence intensities of these regions were obtained using Fiji[5] and the ratios of fluorescence intensity inside the condensate over the fluorescent intensity outside the condensate were calculated. Ratios were only calculated when the region outside the condensate and the region inside the condensate were obtained from the same image frame.

**Fluorescence recovery after photobleaching (FRAP).** Samples were prepared as described for fluorescence microscopy and DIC imaging. Circular regions of interest were irradiated at a laser power of 10% and fluorescence recovery was monitored at 250-millisecond intervals for 30 seconds after bleaching. Image contrast was adjusted globally. The fluorescence intensities for bleached regions were obtained using Fiji[5]. FRAP curves were normalized using a 0 – 1.0 scale by setting the average fluorescent intensities of the four pre-bleach points to be 1.0 and the fluorescent intensity of the first post-bleach point to be 0.

**Transmission electron microscopy.** Aged samples (T ~ 24 hours) from ThT assays were resuspended and removed from the 384 well plates and applied to formvar/carbon coated grids (Electron Microscopy

Sciences, PA). Samples were stained with 2% uranyl acetate and imaged using a JEOL 1400 Transmission electron microscope equipped with an XR80 camera (AMT Imaging, MA) operating at a magnification of 40,000 or 50,000 x.

**Nuclear magnetic resonance spectroscopy.** NMR experiments were conducted on a Bruker Avance NEO spectrometer (Bruker, MA) operating at a proton Larmor frequency of 700.13 MHz at a temperature of 25 °C. Data were processed using NMRPipe[6] or Topspin 4.1.1 (Bruker) and analyzed with CCPNMR Analysis 3.0 software[7]. PU.1 168-270 $^{1}$H, $^{13}$Cα, $^{13}$Cβ, $^{13}$C' and $^{15}$N backbone resonances were assigned using $^{1}$H,$^{15}$N-HSQC, HNCACB, CBCA(CO)NH, HNCO, HN(CA)CO and HCC(CO)NH, recorded on a sample of 425 μM in 50 mM sodium phosphate pH 6.4, 0.5 mM EDTA, 0.02% sodium azide. The $^{1}$H, $^{15}$N-HSQC was recorded using 128* x 1024* complex points in the indirect and direct dimensions, corresponding to acquisition times of 64.4 and 106.5 ms, respectively. The HNCO and HN(CA)CO experiments were recorded using 32* x 32* x 1024* complex points in the indirect (F1, $^{13}$C), (F2, $^{15}$N) and direct (F3, $^{1}$H) dimensions, corresponding to acquisition times of 15.2, 16.1 and 112.6 ms, respectively. The HNCACB and CBCA(CO)NH and HCC(CO)NH experiments were recorded using 100* x 32* x 1024* complex points in the indirect (F1, $^{13}$C), (F2, $^{15}$N) and direct (F3, $^{1}$H) dimensions, corresponding to acquisition times of 8.6, 16.1 and 112.6 ms, respectively. All 3D experiments were recorded using non-uniform sampling (NUS) with a sampling density of 30% for the HNCO and HN(CA)CO experiments and 15% for the HNCACB, CBCA(CO)NH and HCC(CO)NH experiments. Spectra were reconstructed using the SMILE algorithm[8] implemented in NMRPipe.

For the titration of $EWS^{LCD}$ with $^{15}$N labelled PU.1, a ~ 2 mM stock of $EWS^{LCD}$ in 20 mM CAPS pH 11 buffer was titrated into a 550 μL sample containing 50 μM $^{15}$N labelled PU.1 in 50 mM sodium phosphate buffer pH 6.5 to final concentrations of 12.5, 25, 50, 100 and 150 μM. This required the addition of a maximum of 42 μL of $EWS^{LCD}$ stock, thus the concentration of PU.1 is diluted by approximately 7% across the titration. $^{1}$H, $^{15}$N-HSQC spectra were recorded at 25°C for each titration point with 64* x 1024* complex data points in the indirect ($^{15}$N) and direct ($^{1}$H) dimensions, corresponding to acquisition times of 30.1 and 106.5 ms, respectively. Spectra were processed in Topspin 4.1.1 with a sine bell function, zero filled to twice the number of acquired points for data analysis and analyzed using CCPNMR Analysis 3.0 software. Chemical shift perturbations (CSP) were calculated by weighting the $^{1}$H and $^{15}$N chemical shifts with respect to their gyromagnetic ratio using the following equation:

$$\Delta\delta = \sqrt{(\delta\ ^{1}\mathrm{H})^2 + 0.2(\delta^{15}\mathrm{N})^2}$$

CSPs were considered significant when they were higher than the standard deviation of $\Delta\delta_{max}$ for all residues.

$^{15}$N $R_1$ and $R_2$ relaxation rates were calculated from $T_1$ and $T_{1\rho}$ experiments, recorded on a 75 μM sample of PU.1 in 50 mM sodium phosphate pH 6.5, 50 mM NaCl using 64* x 1024* complex data points in the indirect ($^{15}$N) and direct ($^{1}$H) dimensions corresponding to acquisition times of 37.6 and 112.6 ms, respectively. The $^{15}$N $T_1$ experiment consisted of 8 interleaved spectra with the following relaxation delays: 40, 80, 120, 200, 280, 400, 600, and 800 ms. The $T_{1\rho}$ experiment was recorded using a $B_1$ field of 1400 Hz and 8 interleaved spectra with the following relaxation delays: 1, 21, 31, 41, 61, 81, 121 and 161 ms. $^{15}$N $R_2$ rates were calculated using the following equation:

$$R_{1\rho} = R_1\cos^2\theta + R_2\sin^2\theta$$

with $\theta$ = arctan($\omega_1/\Omega$), where $\omega_1$ is the $B_1$ field strength (here 1400 Hz) and $\Omega$ is the offset from the spinlock carrier frequency. $^1$H-$^{15}$N heteronuclear NOE experiments were recorded on the same samples and consisted of two interleaved experiments, with and without proton saturation, using a recycle delay of 4 seconds. Spectra were acquired with 64* x 1024* complex data points in the indirect ($^{15}$N) and direct ($^1$H) dimensions corresponding to acquisition times of 37.6 and 112.6 ms, respectively.

**Electrophoretic mobility shift assays.** 20 µL samples were loaded into wells of a 20% polyacrylamide gel prepared in 0.5 x TBE buffer. Electrophoresis was performed for 60 minutes at 150 V using 0.5 x TBE running buffer. Gels were stained with SYBRsafe (Thermo Scientific, MA) at a 1:20,000 dilution of the DMSO stock in 0.5 x TBE for 30 minutes before imaging using UV transillumination.

**Circular dichroism.** Circular dichroism spectra were recorded using 10 µM samples of all proteins in 50 mM sodium phosphate pH 6.5 at 25 °C in a 2 mm pathlength cuvette using a Jasco 810 spectropolarimeter (Jasco, OK) at a scan speed of 50 nm/min with 0.2 nm increments. Each sample was recorded in triplicate, the data was then averaged and converted to mean residue ellipticity (MRE) using previously described relationships[9]. Melts were conducted by monitoring ellipticity at 220 nm from 25 °C to 95 °C with a data pitch of 0.5 °C and a temperature slope of 1.5 °C/min.

### Supplementary Figures

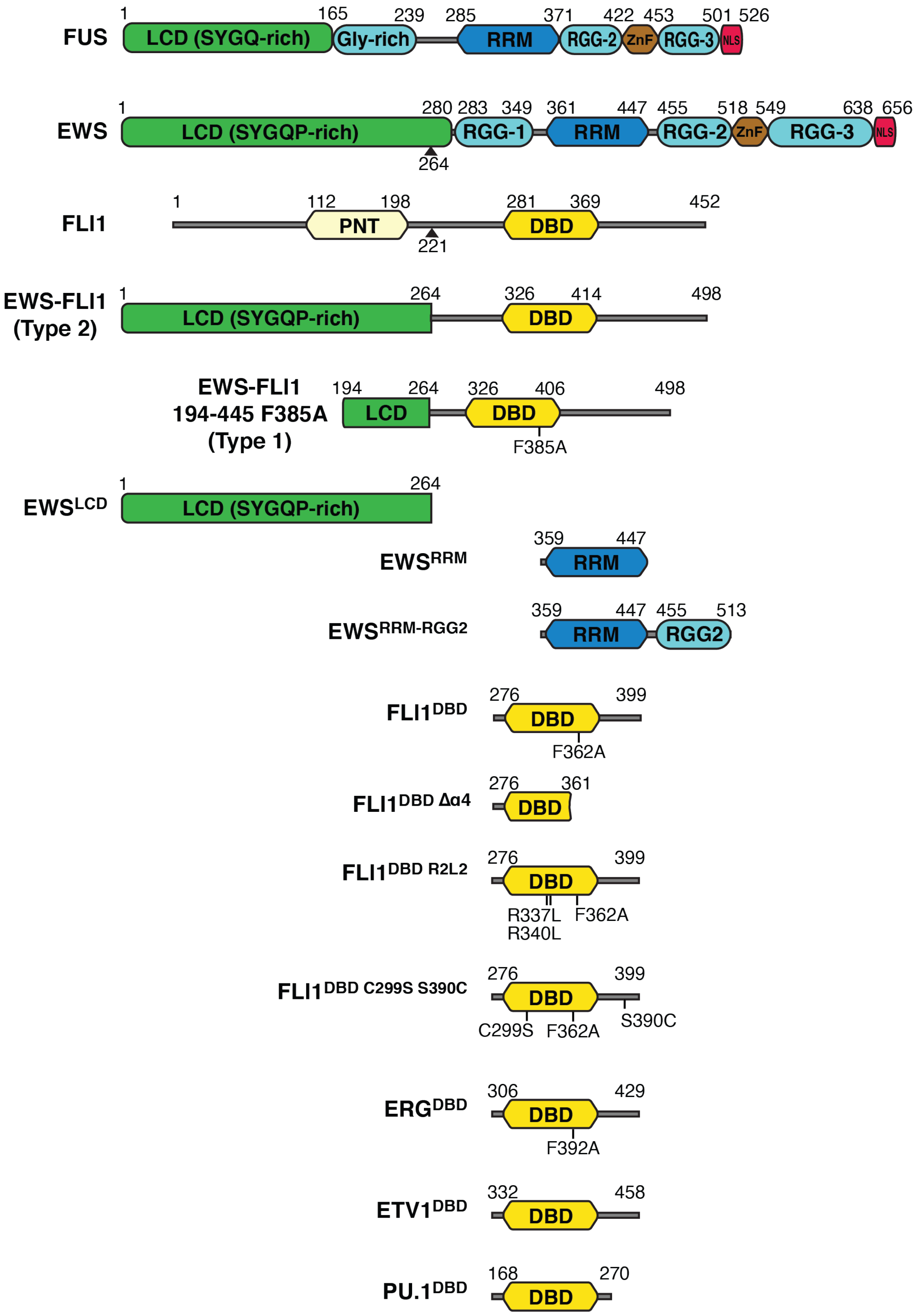


**Supplementary Figure 1. Schematic of the protein constructs used in this study.** Protein sequences and domain margins were extracted from Uniprot and are indicated. Point mutations and fusion break points are indicated with black lines and triangles.

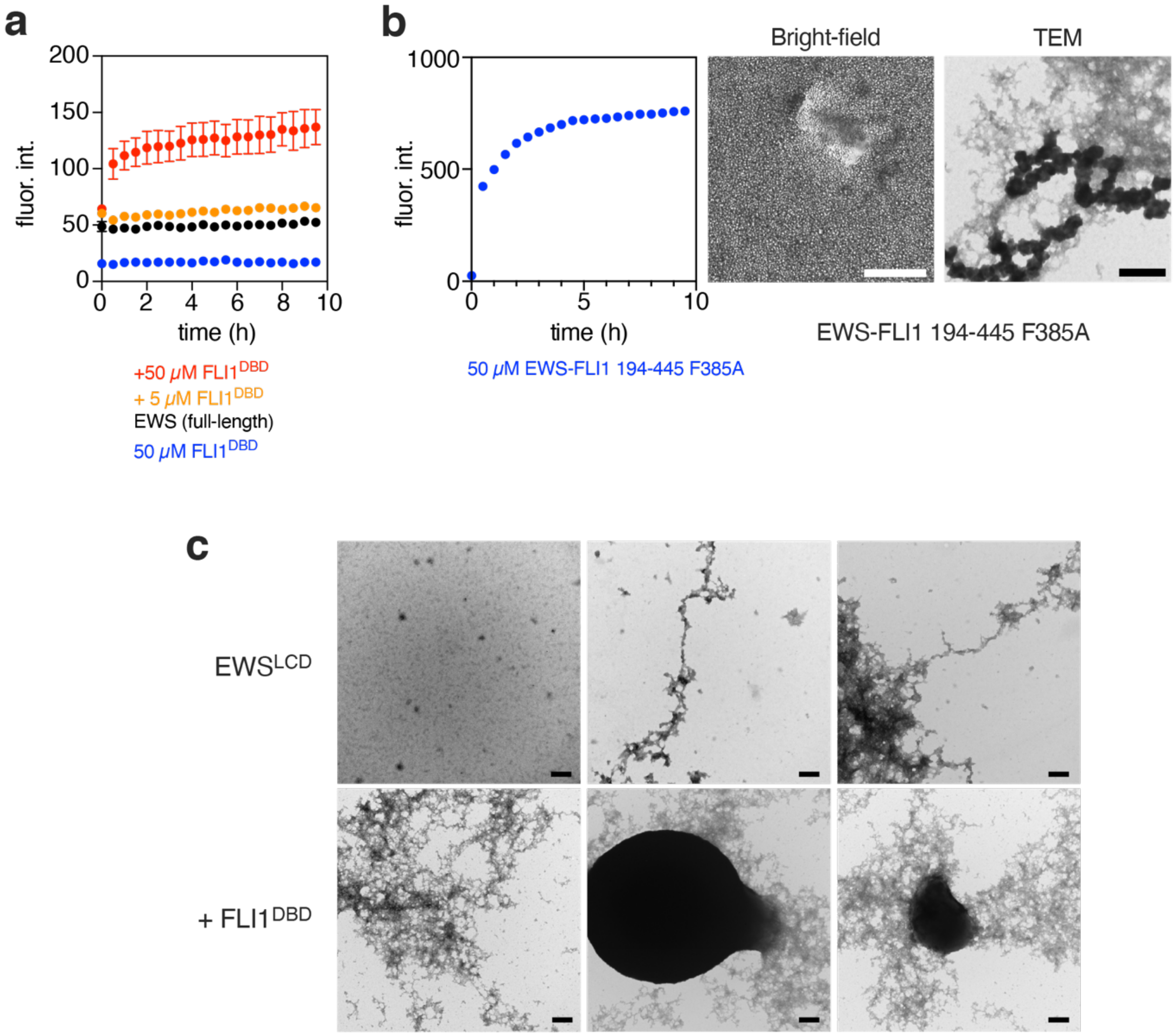


**Supplementary Figure 2. EWS and EWS-FLI1 194-445 F385A form ThT-reactive species. a** ThT assay of 50 µM full-length EWS (black), and with 5 µM FLI1$^{DBD}$ (orange) or 50 µM (red). 50 µM FLI1$^{DBD}$ alone is shown in blue. **b** ThT assay of 50 µM EWS-FLI1 194-445 F385A alone (blue). ThT data are depicted are the average of triplicates ± SEM. Scale bars indicate 50 µm for the bright-field microscopy and 200 nm for the TEM. **c** Representative TEM images of aged samples (T ~ 24 hours) of EWS$^{LCD}$ alone (top panels) and EWS$^{LCD}$ + FLI1$^{DBD}$ (bottom panels). Scale bars indicate 200 nm.

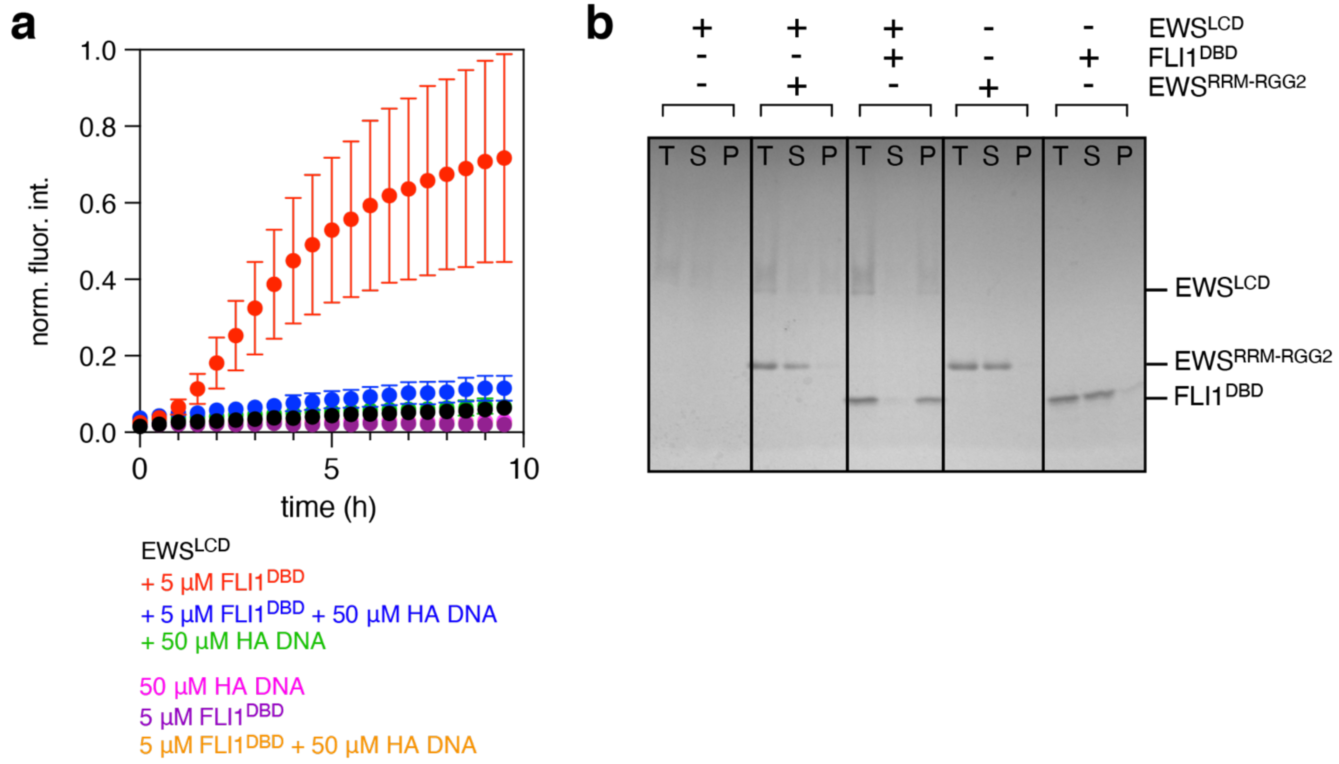


**Supplementary Figure 3. The effect of FLI1$^{DBD}$ on EWS$^{LCD}$ condensates is specific and inhibited by DNA binding. a** ThT assay of 50 µM EWS$^{LCD}$ alone (black) or in the presence of 5 µM FLI1$^{DBD}$ (red), 5 µM FLI1$^{DBD}$ + 50 µM HA DNA (blue) or 50 µM HA DNA (green). Control samples; 5 µM FLI1$^{DBD}$ (purple), 50 µM HA DNA (magenta), and 5 µM FLI1$^{DBD}$ + 50 µM HA DNA (orange). ThT data are depicted are the average of triplicates ± SEM. **b** Pelleting assay of aged samples (T ~ 24 hours) of EWS$^{LCD}$ in 150 mM NaCl, with FLI1$^{DBD}$ or with EWS$^{RRM\text{-}RGG2}$. Control samples; FLI1$^{DBD}$ alone, and EWS$^{RRM\text{-}RGG2}$ alone in 150 mM NaCl. For each sample a lane is shown corresponding to the total (T) sample prior to centrifugation, and the supernatant (S) and pellet (P) after centrifugation.

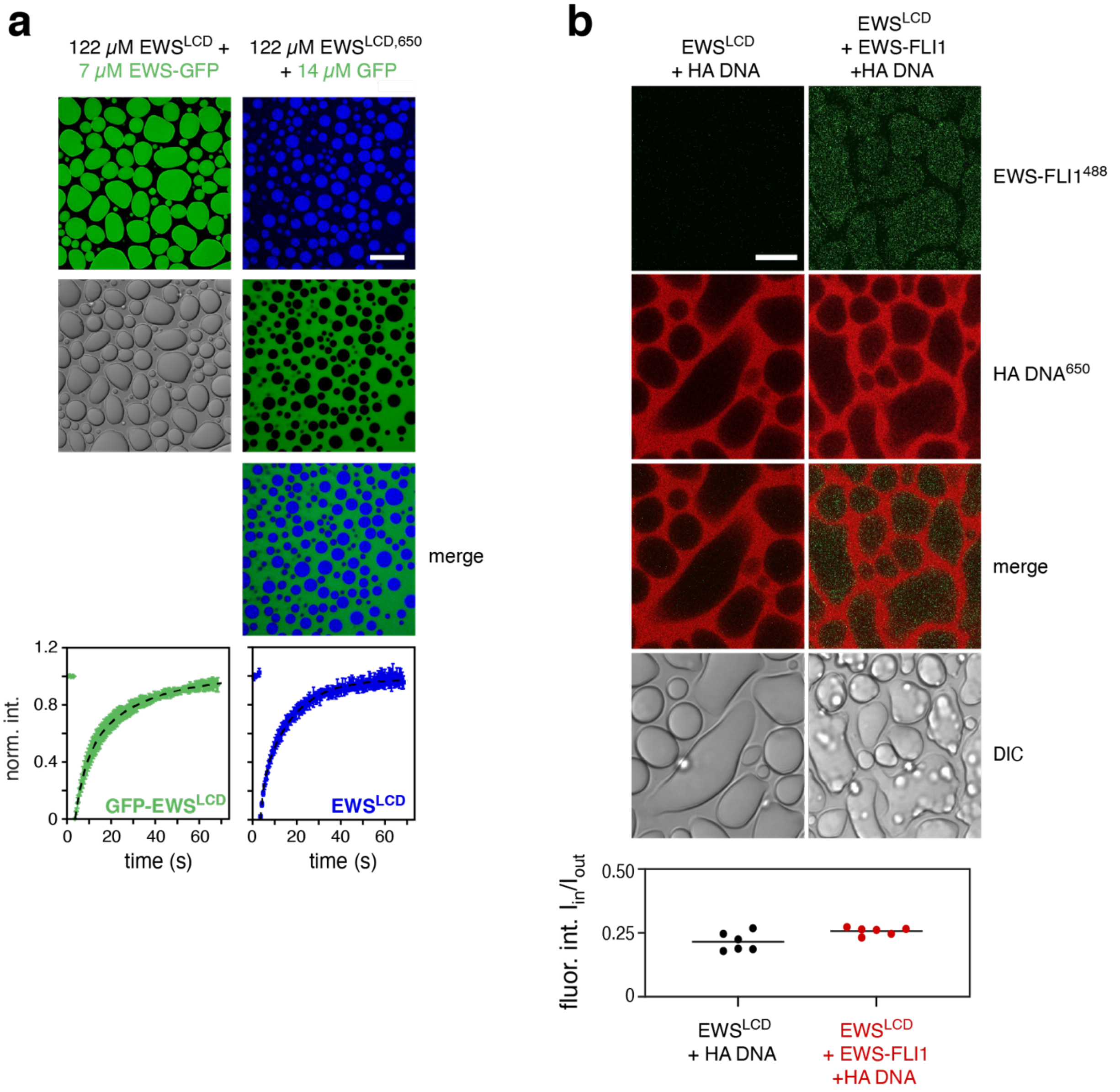


**Supplementary Figure 4. EWS$^{LCD}$ colocalization assays by fluorescence microscopy. a** GFP only colocalizes to EWS$^{LCD}$ condensates when it is fused to EWS$^{LCD}$ (left panels). When EWS$^{LCD\ 650}$ is mixed with free GFP, GFP does not colocalize to the EWS$^{LCD}$ condensates (right panels). Scale bars indicate 25 μm. FRAP analysis of GFP-tagged EWS$^{LCD}$ and EWS$^{LCD\ 650}$ indicates that both constructs form condensates with rapid fluorescence recovery (bottom panels). **b** EWS-FLI1 colocalizes to EWS$^{LCD}$ condensates but HA DNA does not. Unlabeled EWS$^{LCD}$ was mixed with HA DNA$^{650}$ alone (left panels) or with HA DNA$^{650}$ and full-length EWS-FLI1$^{488}$ (right panels). EWS-FLI1 colocalized to condensates containing EWS$^{LCD}$, while HA DNA$^{650}$ was mostly excluded. Scale bars indicate 10 μm. Fluorescence intensity ratios ($I_{in}/I_{out}$) did not indicate a major difference in the degree of colocalization of HA DNA$^{650}$ in the condensates containing EWS$^{LCD}$ and full-length EWS-FLI1 (bottom panel).

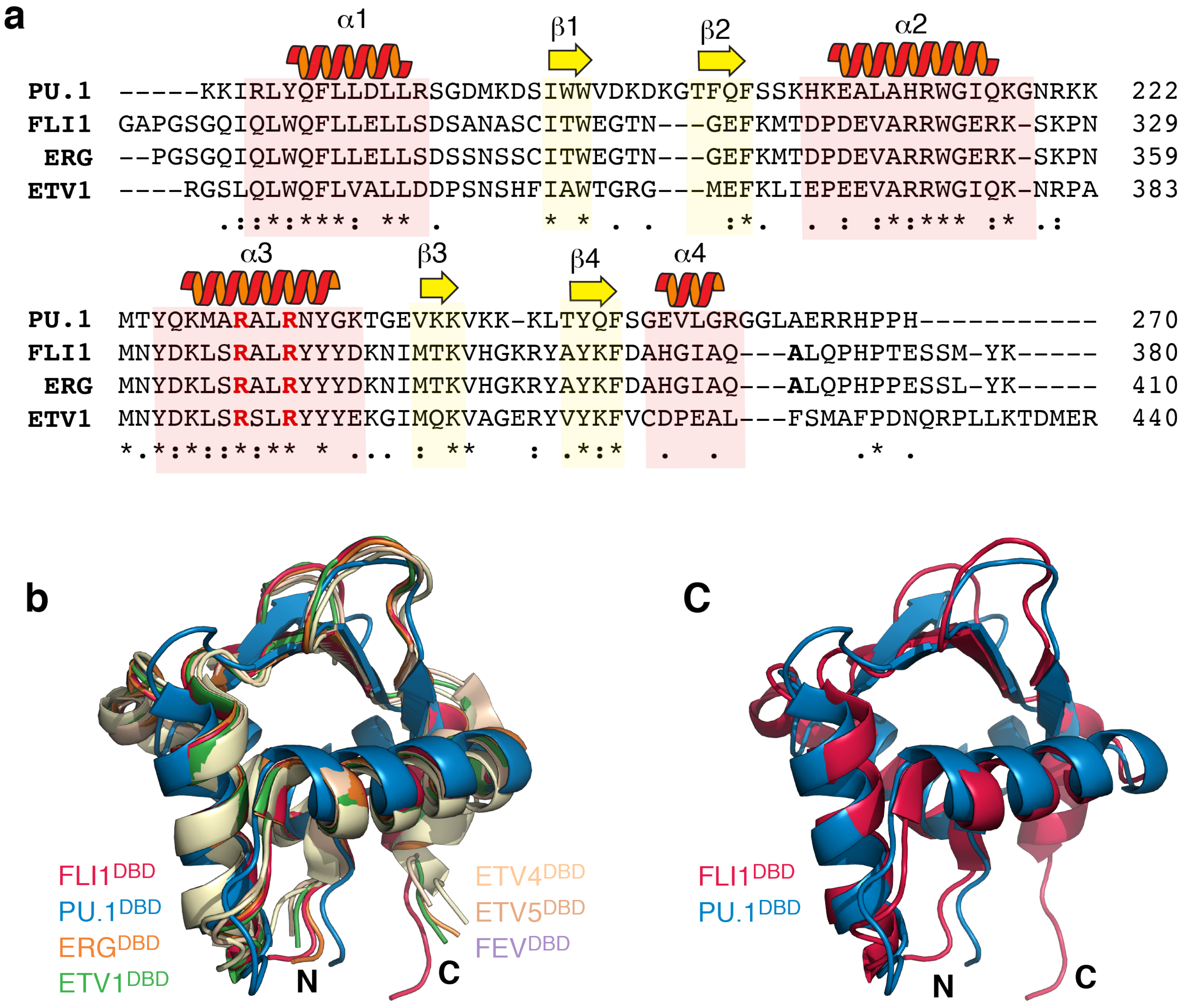


**Supplementary Figure 5. ETS DBDs are highly conserved. a** Amino acid sequence alignment of the DBDs of PU.1, FLI1, ERG and ETV1. Secondary structure assignments are indicated above the sequence and aligned over the sequences with shaded boxes (α-helices, red; β-strands, yellow). Conserved Arg residues involved in DNA binding are bolded in red, F362A (FLI1) and F392A (ERG) mutants are bolded. **b** Overlay of the structures of seven ETS DBDs. **c** Structure overlay of FLI1$^{DBD}$ (red) and PU.1$^{DBD}$ (blue). PDB IDs: FLI1 (5e8g); PU.1 (1pue); ERG (4iri); ETV1 (4bnc); ETV4 (4uuv); ETV5 (4uno); FEV (2ypr).

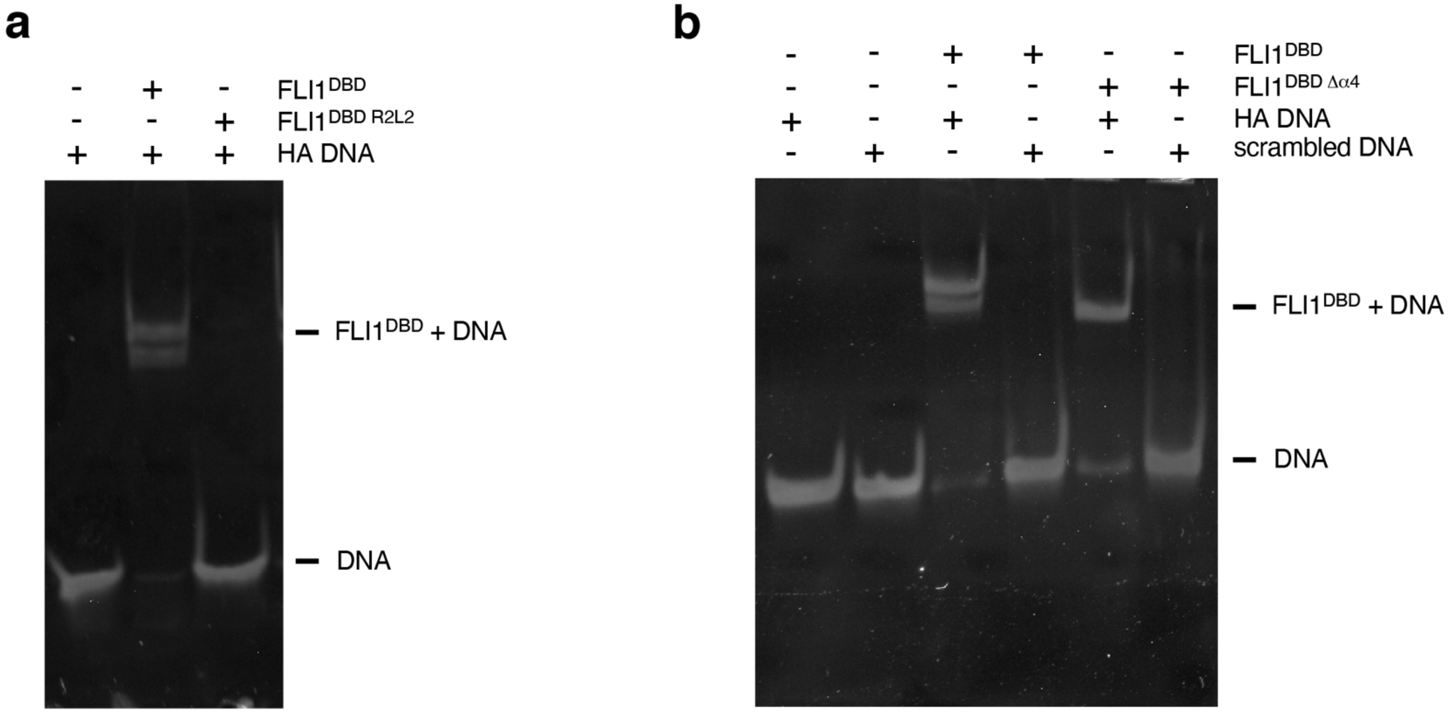


**Supplementary Figure 6. Electrophoretic mobility shift assays of FLI1$^{DBD}$ constructs**. **a** EMSA of FLI1$^{DBD\ R2L2}$ and FLI1$^{DBD}$ with HA DNA. **b** EMSA of FLI1$^{DBD\ \Delta\alpha4}$ and FLI1$^{DBD}$ with HA DNA or scrambled DNA.

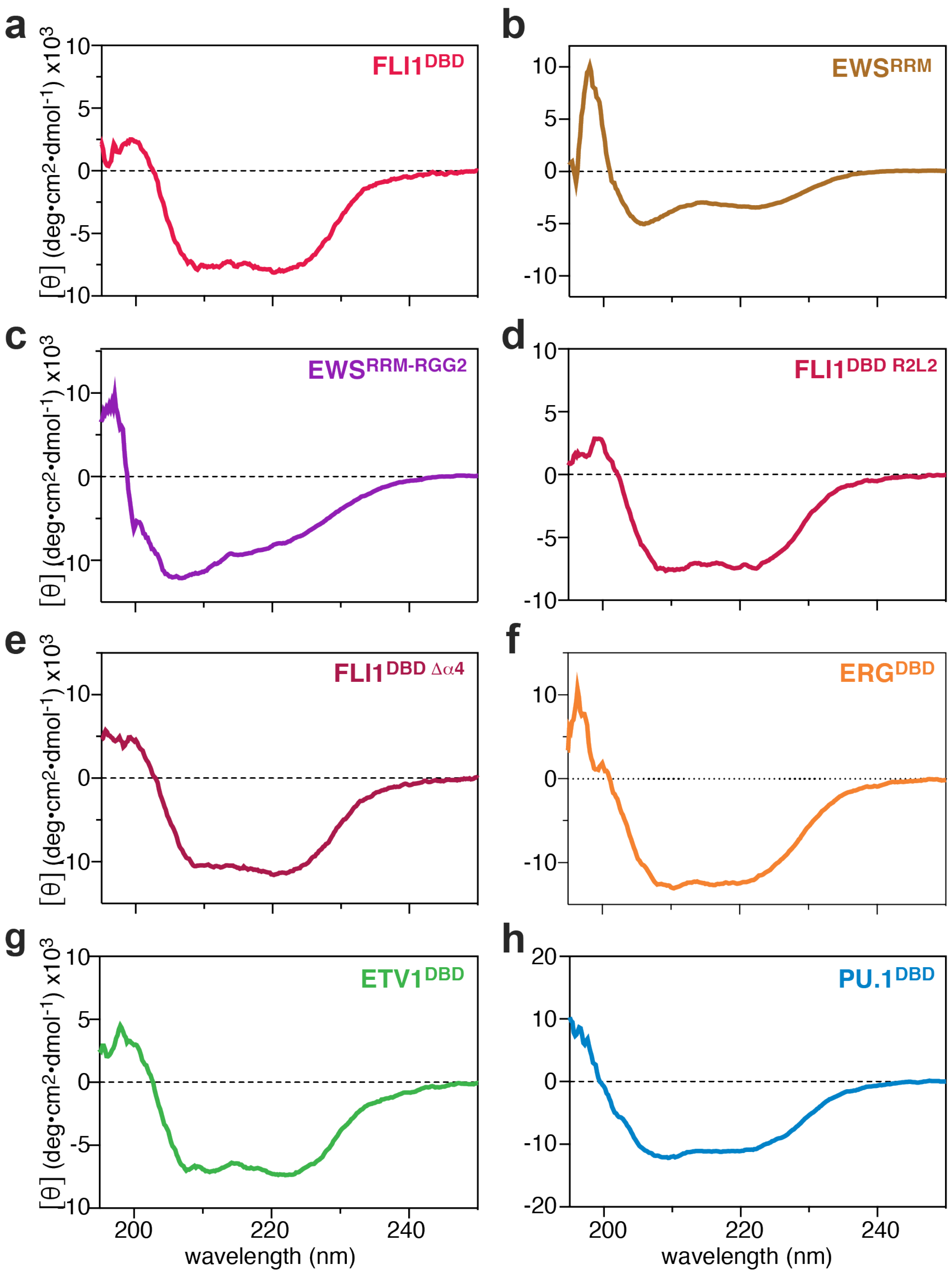


**Supplementary Figure 7. Circular dichroism spectroscopy of ETS DBDs and EWS RNA binding domains.** CD spectra were collected for 10 μM **a** $FLI1^{DBD}$, **b** $EWS^{RRM}$, **c** $EWS^{RRM\text{-}RGG2}$, **d** $FLI1^{DBD\ R2L2}$, **e** $FLI1^{DBD\ \Delta\alpha4}$, **f** $ERG^{DBD}$, **g** $ETV1^{DBD}$, and **h** $PU.1^{DBD}$ at 25 °C in 50 mM sodium phosphate buffer pH 6.5.

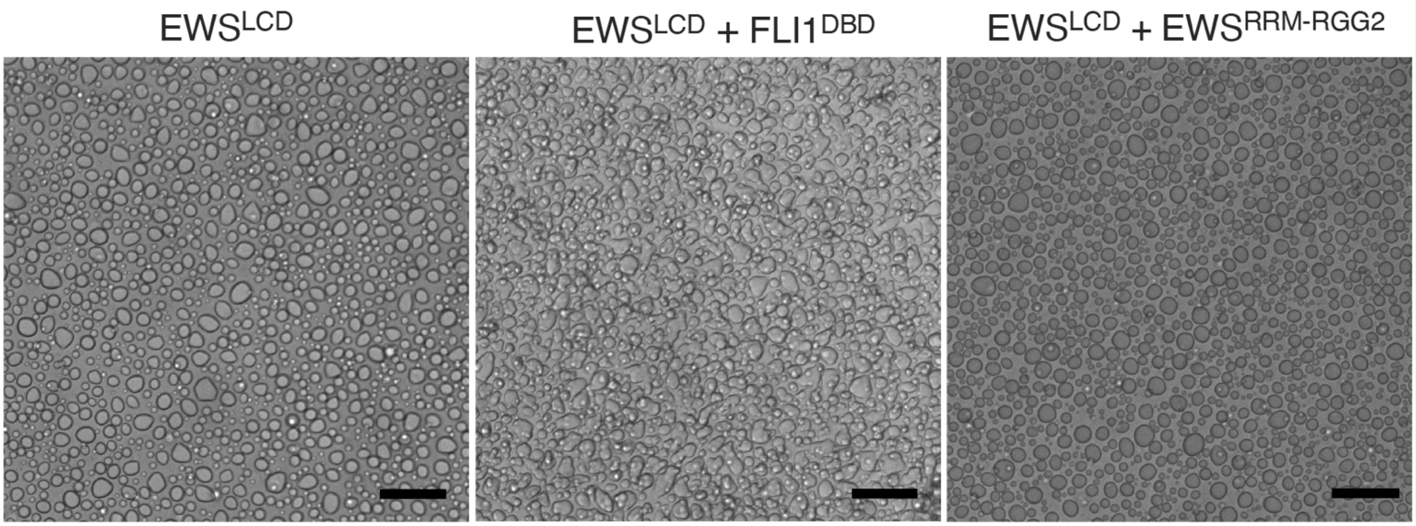


**Supplementary Figure 8. FLI1$^{DBD}$ alters the morphology of EWS$^{LCD}$ condensates.** Bright-field microscopy images taken at T = 0 (right after mixing) of 50 μM EWS$^{LCD}$ alone, or in the presence of 100 μM FLI1$^{DBD}$ or 100 μM EWS$^{RRM-RGG2}$. Scale bars indicate 50 μm.

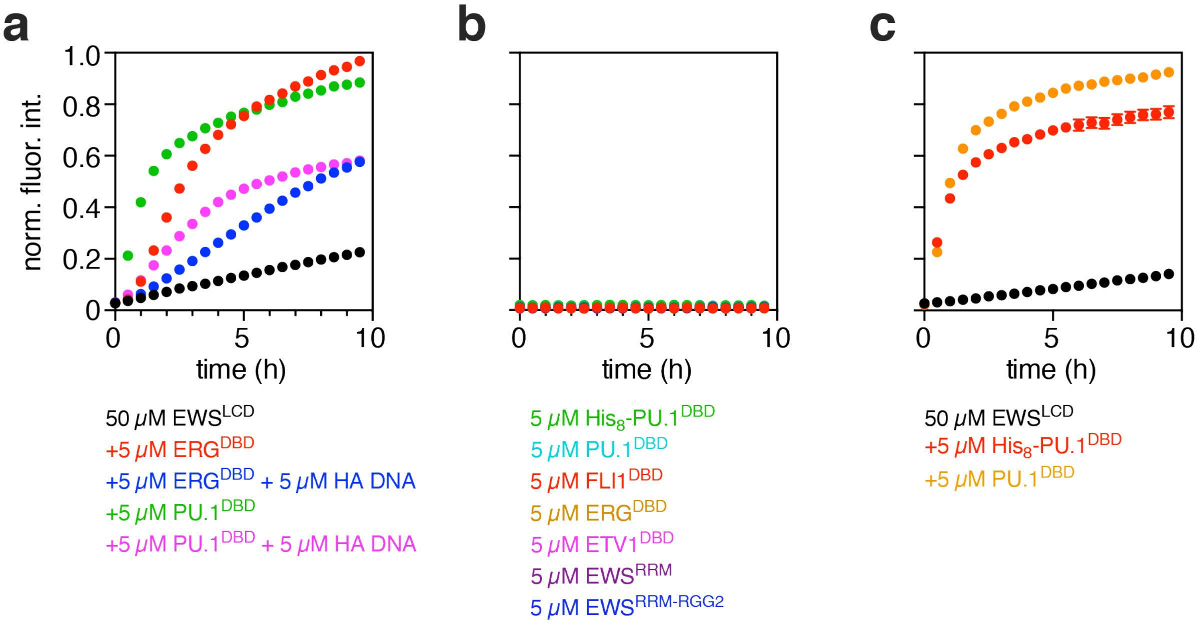


**Supplementary Figure 9. EWS$^{LCD}$ condensate ageing is enhanced by ETS DBDs other than FLI1. a** ThT assays of 50 μM EWS$^{LCD}$ alone (black) or with 5 μM ERG$^{DBD}$ (red), 5 μM ERG$^{DBD}$ + 5 μM HA DNA (blue), 5 μM PU.1$^{DBD}$ (green) or 5 μM PU.1$^{DBD}$ + 5 μM HA DNA (magenta). **b** Control ThT assays of all ETS DBDs and EWS RNA-binding domain constructs used in this study incubated alone at 5 μM as indicated. **c** ThT assays of 50 μM EWS$^{LCD}$ alone (black) or with 5 μM His$_8$-tagged PU.1$^{DBD}$ (red) or 5 μM PU.1$^{DBD}$ (orange). ThT data is presented as the average of triplicates ± SEM.

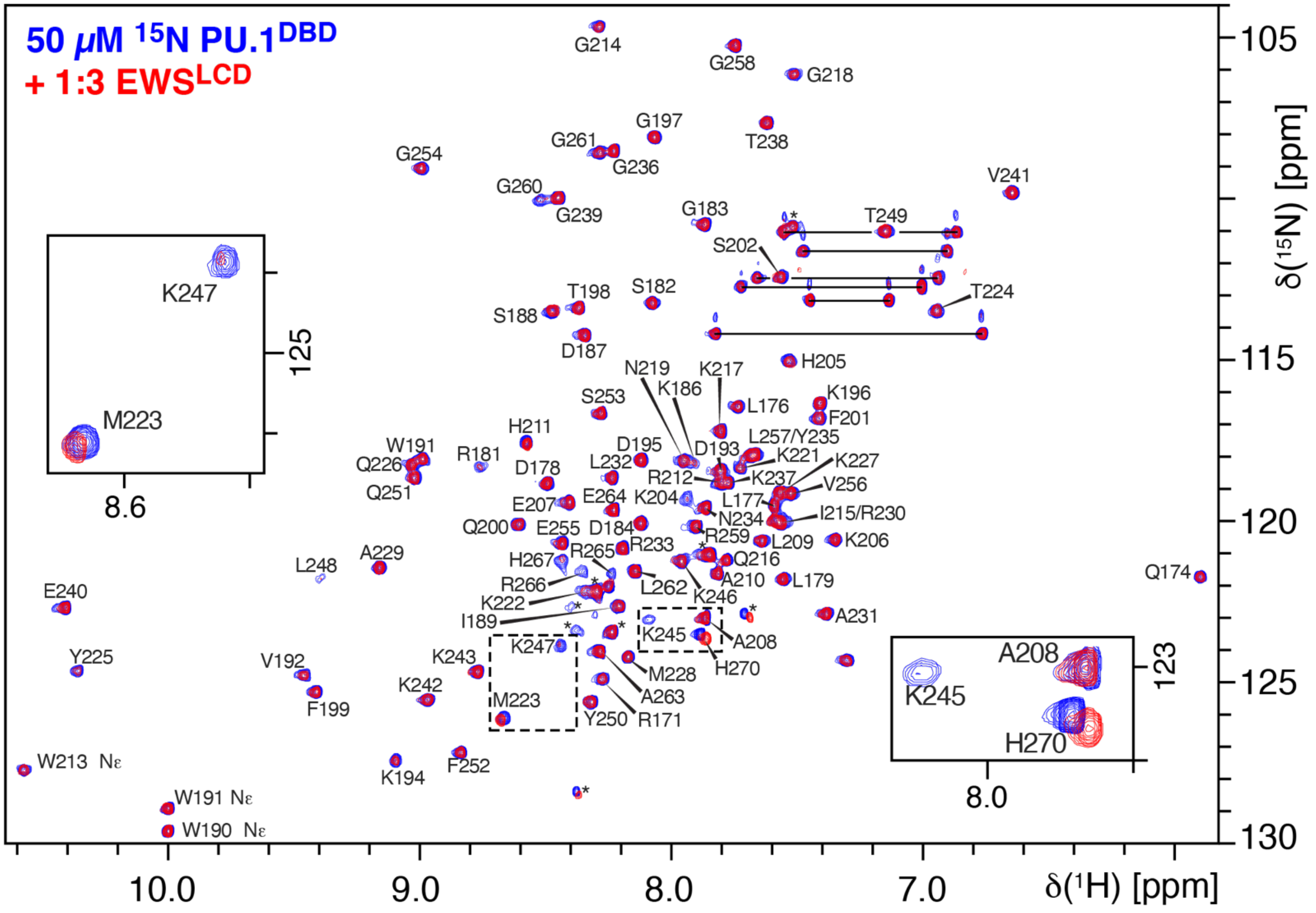


**Supplementary Figure 10. Titration of PU.1$^{DBD}$ with EWS$^{LCD}$.** $^{1}$H,$^{15}$N-HSQC of 50 μM PU.1$^{DBD}$ (blue) and with a 3:1 molar ratio of EWS$^{LCD}$ (red). Peaks are labeled using the one-letter amino acid code. Insets show regions in which peaks are either broadened or shifted because of EWS$^{LCD}$ interactions. Resonance assignments are deposited in the BMRB (XXXXX).

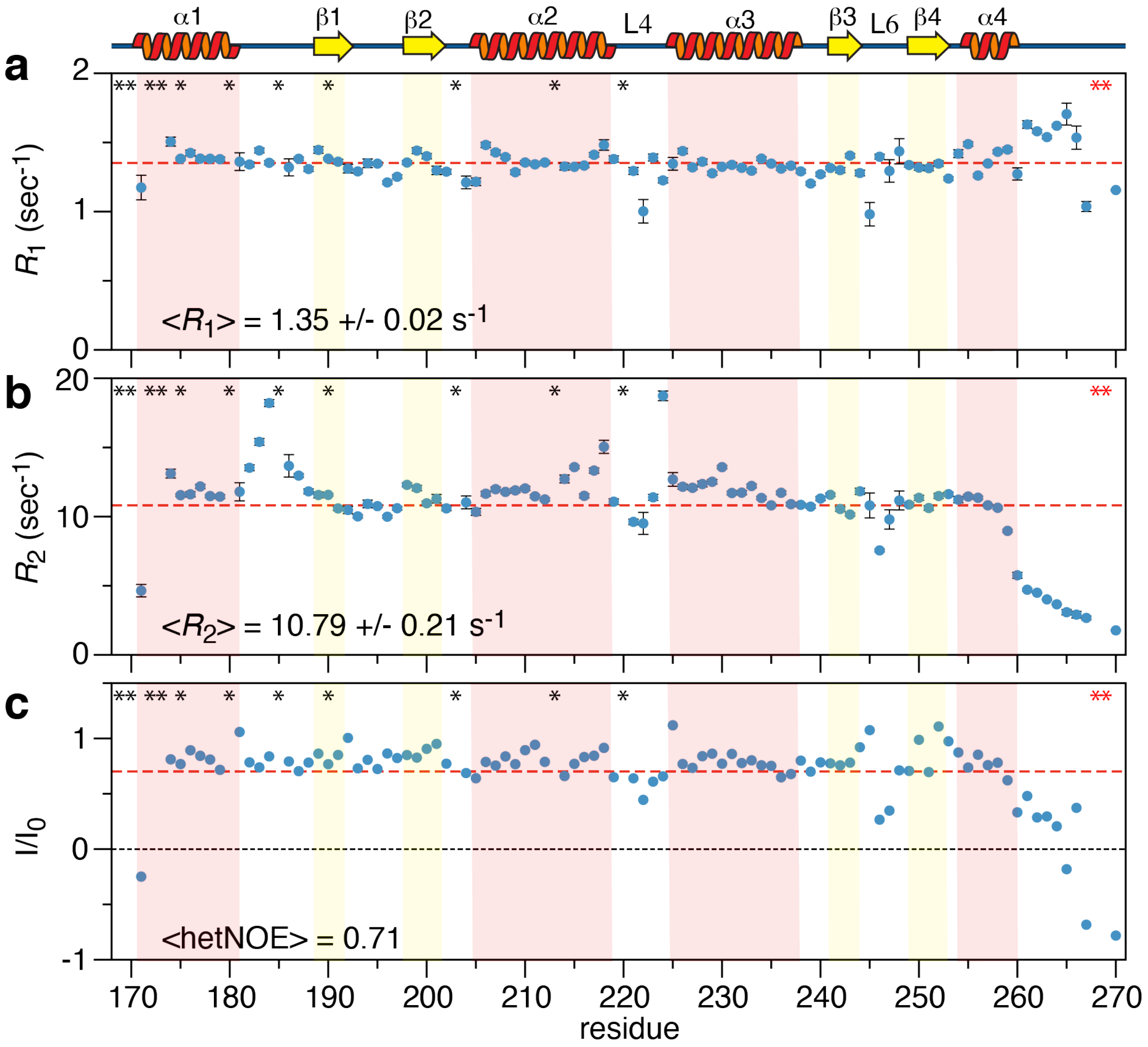


**Supplementary Figure 11. Nuclear spin relaxation parameters for PU.1$^{DBD}$**. **a** $R_1$, **b** $R_2$, and **c** hetNOE were measured on a 75 μM sample of PU.1$^{DBD}$. Secondary structure elements from the crystal structure (PDB 1pue) and are depicted in cartoon format and aligned across all panels with shaded boxes. In each of **a**, **b**, and **c,** dashed red lines indicate the average value and standard deviation of $R_1$, $R_2$ or hetNOE. Black asterisks denote residues that are overlapped or unassigned. Red asterisks indicate the two proline residues.

### References


1. Monahan, Z. et al. Phosphorylation of the FUS low-complexity domain disrupts phase separation, aggregation, and toxicity. *EMBO J* **36**, 2951-2967 (2017).
2. Hou, C. & Tsodikov, O.V. Structural Basis for Dimerization and DNA Binding of Transcription Factor FLI1. *Biochemistry* **54**, 7365-74 (2015).
3. Johnson, C.N., Xu, X., Holloway, S.P. & Libich, D.S. The (1)H, (15)N and (13)C resonance assignments of the low-complexity domain from the oncogenic fusion protein EWS-FLI1. *Biomol NMR Assign* **16**, 67-73 (2022).
4. Antos, J.M. et al. Site-Specific Protein Labeling via Sortase-Mediated Transpeptidation. *Curr Protoc Protein Sci* **89**, 15 3 1-15 3 19 (2017).
5. Schindelin, J. et al. Fiji: an open-source platform for biological-image analysis. *Nat Methods* **9**, 676-82 (2012).
6. Delaglio, F. et al. NMRPipe: a multidimensional spectral processing system based on UNIX pipes. *J Biomol NMR* **6**, 277-93 (1995).
7. Skinner, S.P. et al. CcpNmr AnalysisAssign: a flexible platform for integrated NMR analysis. *J Biomol NMR* **66**, 111-124 (2016).
8. Ying, J., Delaglio, F., Torchia, D.A. & Bax, A. Sparse multidimensional iterative lineshape-enhanced (SMILE) reconstruction of both non-uniformly sampled and conventional NMR data. *J Biomol NMR* **68**, 101-118 (2017).
9. Myers, J.K., Pace, C.N. & Scholtz, J.M. Helix propensities are identical in proteins and peptides. *Biochemistry* **36**, 10923-9 (1997).
